# Taurine Inhibits Apolipoprotein E4 Aggregation

**DOI:** 10.1101/2025.08.13.669519

**Authors:** Anthony Legrand, Katerina Amruz Cerna, Sérgio M. Marques, Naina Verma, Jakub Kopko, Tereza Vanova, Madhumalar Subramanian, Jaroslav Bendl, Tomas Henek, Pavel Vanacek, Josef Kucera, Joan Planas-Iglesias, Jiri Sedmik, Veronika Pospisilova, Petr Kouba, Aneta Vaskova, Marketa Nezvedova, Jiri Sedlar, Jiri Damborsky, Stanislav Mazurenko, Martin Marek, Josef Sivic, Lenka Hernychova, David Bednar, Dasa Bohaciakova, Zbynek Prokop

## Abstract

Apolipoprotein E4 (ApoE4) is a major genetic risk factor in many neurodegenerative diseases, yet effective therapeutic strategies targeting its associated pathologies remain unresolved. The aggregation of ApoE4, a key pathological feature, can be attenuated by tramiprosate and its metabolite 3-sulfopropanoic acid. In this study, we investigated the potential of taurine, a close chemical analogue of tramiprosate, to modulate ApoE4-mediated pathological processes. Using an integrated approach—including molecular dynamics simulations, static light scattering, mass spectrometry, and cerebral organoid models—we investigated taurine’s effects on ApoE4 aggregation. We found that taurine effectively inhibits ApoE4 aggregation. Notably, taurine significantly ameliorates the pathophysiological characteristics of ApoE4, bringing its phenotype closer to the more benign ApoE3 variant. By leveraging its neuroprotective properties, taurine may offer effects comparable to tramiprosate and 3-sulfopropanoic acid, positioning it as an accessible and promising candidate for mitigating neurodegeneration, particularly in individuals with the high-risk *ApoE4/E4 genotype*.

## Background

Human apolipoprotein E (ApoE) is a 34 kDa glycoprotein that was initially isolated as a very low-density lipoprotein (VLDL) from plasma. It is structured into an N-terminal 5-helix bundle (**Figure 1A**), followed by a hinge region and a C-terminal membrane-anchoring domain.^1^ ApoE is involved in lipid metabolism as well as a variety of neurodegenerative diseases, such as macular degeneration, Alzheimer’s Disease (AD), Parkinson’s disease (PD), Lewy body dementia (LBD), and macular degeneration.^2,3^ ApoE is the key molecule for pathogenesis at various levels of biological systems, formulated in the so-called “ApoE Cascade Hypothesis”. This hypothesis states that the biochemical and biophysical properties of ApoE impact a cascade of events at the cellular and systems levels, ultimately impacting aging-related pathogenic conditions, such as AD. However, such involvement vastly depends on which isoform is encoded in the patient’s genome. In the case of AD and PD, ApoE2 is a protective factor, ApoE3 is neutral, and ApoE4 is a risk factor.^4^ Previously, we described and compared ApoE3 and ApoE4 self-associating interfaces (**Figure 1B**), and how tramiprosate (TMP) and its metabolite 3-sulfopropanoic acid (SPA)^5^ can inhibit their formation (**Figure 1C**). We highlighted the domino-like effect of the single mutation from ApoE3 to ApoE4, C112R, leading to multiple conformational changes propagating through the entire protein and modifying aggregation-prone interfaces, which could be then corrected using SPA. Additionally, we showed that COs could endogenously metabolise TMP into SPA, and that such treatment stimulates neurogenesis, modifies cholesterol metabolism and the levels of AD-related proteins, and decreases MAP kinase signaling in cerebral organoids (COs) with ApoE4/E4 genotype.

**Figure 1.**
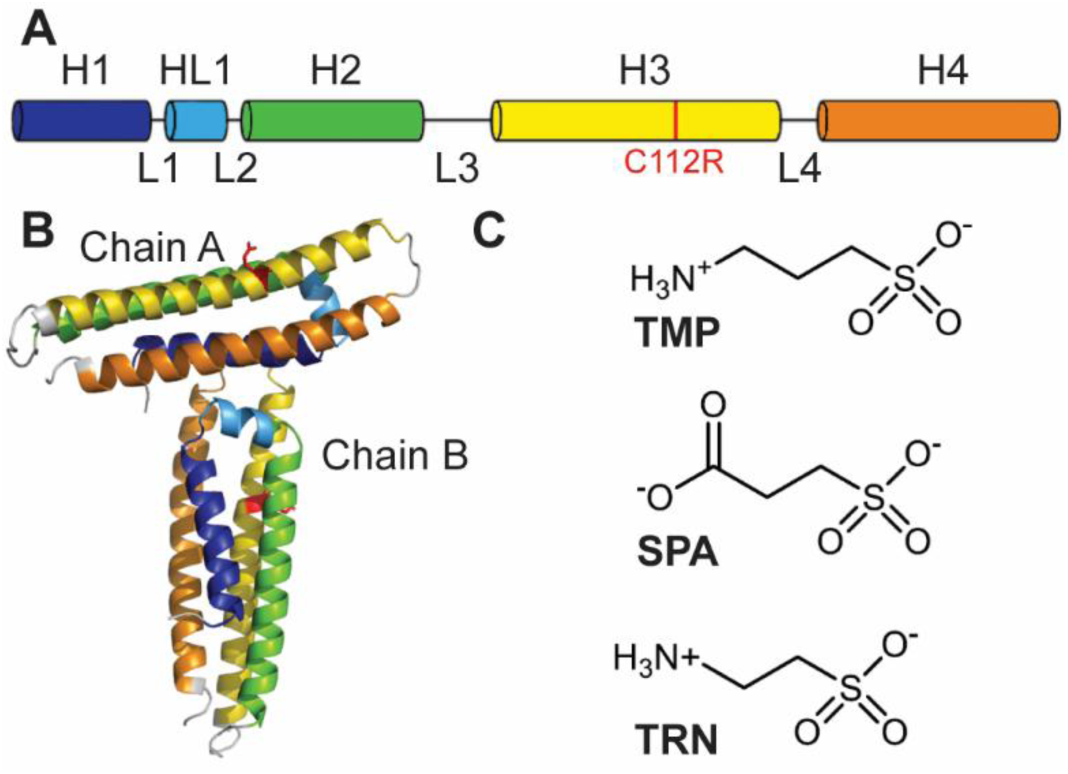
Description of ApoE’s N-terminal domain and small molecules used in this study. (A) The N-terminal domain of ApoE4 (23-166) is a 5-helix bundle. C112R mutation is highlighted in red. H: helix. HL1: small helix between H1 and H2. L: loop. (B) Structure of an ApoE4_23-166_ T-shaped dimer assembled from 2 monomers (PDB: 8AX8). The rest of the protein was not resolved experimentally and is thus absent from the final structure. Through this study, we refer to chains A and B as the horizontal and vertical protomers, respectively. (C) Molecular structure of tramiprosate (TMP), 3-sulfopropanoic acid (SPA), and taurine (TRN).

TRN is a highly abundant brain metabolite, mainly detected in the central nervous system (**Figure 1C**). Its supplementation, usually through diet, is beneficial in the contexts of tissue and cell stress, inflammation and in many pathologies, such as intracerebral haemorrhage or hypertension.^6^ TRN has a very similar structure to TMP, only differing by a single methylene group. While TMP is metabolized *in vivo* into SPA,^5^ TRN is obtained from cysteine and metabolized into dipeptides (e.g. glutamyltaurine), acyltaurines, and TRN-conjugated bile acids.^7^ Finally, there is a clear correlation between decreased TRN levels in the brain, brain senescence, and the accumulation of phosphorylated protein Tau and Amyloid-beta (Aꞵ) peptide.^8^ However, to the best of our knowledge, its putative interplay with ApoE in the context of neurodegeneration has never been addressed. Here, we combine *in silico, in vitro*, and *in vivo* cellular experiments to characterize the influence of TRN on ApoE aggregation and to evaluate its effects on cerebral organoids.

## Methods

### System preparation and equilibration

The structure of TRN was constructed and minimized using Avogadro 2.^9^ As expected at pH 7.4, SPA was considered deprotonated on the sulfonate (SO^3-^) and protonated on the amine (NH^3+^) groups, with a net charge of 0. A minimization step was performed on those structures by the Auto Optimization Tool of Avogadro, using the UFF force field with the steepest descent algorithm.^10^ The resulting structure was then submitted for further optimization and calculation of their partial atomic charges using Gaussian 09,^11^ with the Hartree-Fock method and 6-31G(d) basis set in vacuum. The antechamber module of AmberTools 16^12^ was used to extract the RESP charges of the ligand from the Gaussian output files. To prepare the PAR and RTF parameter files compatible with the CHARMM force field we used the SwissParam web server.^13^

The three-dimensional structures of the N-terminal domain (23-166) of ApoE3 and ApoE4 were obtained from the RCSB Protein Data Bank (PDB ID 1BZ4 and 8AX8, respectively).^5,14^ PyMOL 2.3 (The PyMOL Molecular Graphics System, Version 2.3 Schrödinger, LLC.) was used to generate the T-shaped dimer from that structure with the symmetry information contained in the original PDB files. Both protomer chains of the dimers were renamed as A and B (Fig. 1B). The crystallographic water molecules were maintained.

The following steps were performed with the High Throughput Molecular Dynamics (HTMD) scripts.^15^ The protein was protonated with PROPKA 2.0 at pH 7.4.^16^ Each protein was simulated free or with an excess of TRN ligands. The free protein was solvated with a cubic box of TIP3P water with the edges at least 25 Å away from the protein atoms, using the solvate module of HTMD.^17^ Using a Python script in the preparation protocol of HTMD, the protein with ligands was prepared by adding 100 molecules of TRN randomly placed on a sphere with a radius of 3 Å further from any atom of the protein, and at least 5 Å away from each other; the system was then solvated with a cubic box of TIP3P water with the edges at least 5 Å away from the ligand atoms. Na^+^ and Cl^-^ ions were added to neutralize the charge of the system and get a final salt concentration of 0.1 M. The topology of the system was built using the htmd.builder.charmm module of HTMD, with the modified CHARMM36m^18^ force field and the parameters for the modified mTIP3P^17^ solvent model and the respective parameters for the ligand.

Each system was equilibrated using the Equilibration_v2 module of HTMD.^15^ It was first minimized using the conjugate-gradient method for 500 steps. Then, the system was heated to 310 K and minimized as follows: (i) 500 steps (2 ps) of NPT thermalization with the Berendsen barostat with 1 kcal·mol^-1^·Å^-2^ constraints on all heavy atoms of the protein, (ii) 1,250,000 steps (5 ns) of NVT equilibration with Langevin thermostat and the same constraints, and (iii) 1,250,000 steps (5 ns) of NVT equilibration with the Langevin thermostat without any constraints. During the equilibration simulations, holonomic constraints were applied on all hydrogen-heavy atom bond terms and the mass of the hydrogen atoms was scaled with factor 4, enabling a 4 fs time step.^19–22^ The simulations employed periodic boundary conditions, using the particle mesh Ewald method for the treatment of interactions beyond a 9 Å cut-off, and the smoothing and switching of van der Waals interaction was performed for a cut-off at 7.5 Å.

### Adaptive sampling MDs to study dissociation

HTMD was used to perform adaptive sampling molecular dynamics simulations (MDs) to study the dissociation process of the ApoE dimers. For that, 50 ns production MD runs were started with the systems that resulted from the equilibration cycle and employed the same settings as the last step of the equilibration. The trajectories were saved every 0.1 ns. Adaptive sampling was performed using the root-mean square deviation (RMSD) of the Cɑ atoms with respect to the initial structure, and time-lagged independent component analysis (tICA) in 1 dimension.^23^ Several epochs of 10 MDs each were performed for each system, until the dissociation of the dimers was observed or it reached a maximum of 20 epochs (corresponding to a cumulative time of 10 µs).

### Adaptive goal MDs to study self-association

We performed an adaptive sampling of the ApoE with the adaptive goal method implemented in HTMD to study the association process of the two chains into a dimer. For that, we identified, for each system, the individual MD from the previous adaptive sampling that displayed the most dissociated state of the dimer (identified by the highest RMSD of the Cα atoms). This MD was used as a starting point for the adaptive goal simulations.

For ApoE4 without SPA or TRN, the dissociation was not observed before 10 μs. In this case, the two chains in the initial dimer were manually split, and with the HTMD scripts, they were randomly placed 15 Å apart from each other. The resulting system was solvated with a cubic box of TIP3P waters with the edges 22.5 Å from any atoms of the protein, and Na^+^ and Cl^-^ ions were added to neutralize the system and achieve 0.1 M concentration of salt. This system was further parametrized and equilibrated as described above. This ensured a system of nearly the same size as for the previous adaptive sampling.

Adaptive goal simulations were performed with the goal function defined to minimize the RMSD of the Cα atoms with respect to the initial crystal structure of the dimer (in a T-shape conformation). The exploration/exploitation ratio of the adaptive method (adg.ucscale) was set to 0, to favor only the exploitation and disregard exploration. In doing so, we encouraged the systems to sample structures closer and closer to the T-shaped dimers. The MDs were run with the same settings as the last step of the equilibration described above. Overall, 19 or 20 epochs of 10 MDs each were performed for each system, corresponding to a cumulative time of ca. 10 µs.

### Post-processing of molecular dynamics simulations

The previous adaptive sampling and adaptive goal simulations were converted into a simulation list using HTMD,^15^ the water and ions were filtered out, the system was wrapped (to place all the molecules back into a single periodic simulation box), and unsuccessful MDs shorter than 50 ns were omitted. ParmEd was used to convert the CHARMM topologies to AMBER topologies.^24^ The cpptraj^25^ module of AmberTools 16 (Case et al., 2016) was used to concatenate the trajectories of each system, ordering the epochs chronologically, center and align them by their Cα atoms, and save the combined trajectory in a single file for further analysis.

### Interaction energies

The linear interaction energy (LIE)^26^ was computed to assess the binding energy of each residue in the dimers of ApoE3 and ApoE4 with the SPA or TRN molecules, expressed as the respective electrostatic and van der Waals components, in the respective adaptive simulations. To this end, we used cpptraj to calculate the LIE on every snapshot of the combined MD trajectories. These interactions were calculated when the dimers were still associated, while we considered only the first 500 ns of the concatenated MD. This method was also used to assess the interaction energy between the two chains of the ApoE dimers in every system, with and without small molecules. These calculations were also performed on the first 500 ns of the concatenated adaptive simulations.

### Dimensionality reduction and clustering analysis

The association adaptive goal simulations were analyzed by dimensionality reduction with the variational approach for Markov processes (VAMP) (Wu and Noé, 2019), a technique suitable for detecting metastable states in off-equilibrium settings.^27^ It is a method based on the Koopman theory, projecting the data onto the dominant components of the Koopman operator, which represent the slowest estimated processes. To ensure an unbiased exploration of all conformations formed during the simulation, all inter-chain inter-residue distances were used as input. A lag time of 15 ns was used. The data was projected onto the first 5 dominant VAMP dimensions. Frames with a minimum inter-heavy atom distance between the two ApoE chains exceeding 8 Å were filtered out and used to calculate the fraction of monomeric state in each system. After filtering, K-means clustering was applied to identify the metastable states.

### Analysis of RMSD from reference structures

To further validate the results of VAMP clustering and provide a fair comparison between different systems, the RMSD to the reference structures of interest was computed for all frames across all ApoE3 and ApoE4 simulations (with and without TRN and SPA). The crystallographic T-shaped dimer, V-shaped dimer, parallel dimer, and the new conformation observed in our simulations, named anti-T-shaped dimer, were used as the reference structures. The reference structures of the T-shaped, V-shaped (previously described in ^5^) and parallel dimers were obtained from the respective crystallographic structures by retrieving the symmetry mates with PyMOL, while the anti-T-shaped dimer reference structure was determined by selecting a frame from the respective cluster with strong LIE and visual assessment. The RMSD value was computed for both possible alignments of the two chains, i.e., A and B, taking the minimum value for each frame. The number of frames with RMSD to a given reference structure lower than the threshold of 6 Å was determined. This method allows obtaining a numerical value determining the proportion of frames close to the given reference structure for each system (even if the system lacked a VAMP cluster representing the given reference structure) and ensures that the discussed populations represent frames closely resembling reference structures.

### Static light scattering (SLS)

SLS was recorded at 266 nm using an UNcle screening platform (UNchained Labs) at 37 °C. To this end, 10 µM of commercially available full-length ApoE3 or ApoE4 (PeproTech) in 10 mM Tris 50 mM NaCl pH=7.4 was mixed with 20 mM SPA, 20 mM TRN, or buffer (drug stock solutions were at 0.5 M in 1M Tris pH=7.4) and then monitored for 14 h. Experiments were performed with three technical triplicates. Analytical fits were performed with KinTek (KinTek Corporation), with a single exponential function with linear drift component (*Eq. 1*), with *A*_0_ as a constant baseline, *A*_1_ and *k*_obs,1_ as amplitude and observed rate for the initial exponential phase, *k*_obs,2_ as observed rate of follow-up linear drift:

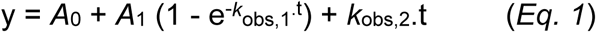

We computed the change in aggregation as the drop in *A*_1_ between ApoE4 alone (*A*_1_(ApoE4)) and in the presence of drug (*A*_1_(drug)) (*Eq. 2*):

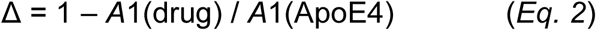

T-test is performed using NORM.DIST function on Excel 2023 (Microsoft).

### HDX-MS

Firstly, undeuterated samples of ApoE3 and ApoE4 proteins were prepared and analyzed separately using a mass spectrometer operated in data-dependent MS/MS mode with PASEF enabled. The identified peptides provided information regarding their satisfactory number, sequence coverage, and redundancy. These optimized sample preparation and analysis conditions were then applied to HDX-MS analyses.

Control undeuterated samples of ApoE3/4, both free and in complex with SPA or TRN, were diluted to a final concentration of 2 µM with a basic H2O buffer (10 mM Tris-HCl, 50 mM NaCl, pH 7.5). A deuterated buffer of identical composition (pD 7.5, pH 7.1) was used for preparing deuterated samples. The Leap HDX robotic station (Trajan Scientific and Medical) was used to prepare each HDX time point by mixing 3 μl of protein, 3 μl of ligand in a 1:1000 molar ratio followed by pre-incubation at 2°C for 5 min and then dilution with 59 uL D2O buffer making the final protein concentration to 2 µM, and the D2O buffer composition was 90%. Four different time points, 60 s, 120 s, 600 s, and 1800 s, were quenched by adding a quenching buffer maintained at 2 °C (6 M urea in 1 M glycine, pH 2.3) in a 1:1 ratio. All samples were prepared in triplicates.

100 pmoles of each quenched sample were directly injected into an immobilized dual pepsin column (Pepsin/Nepenthesin2, AffiPro) at a flow rate of 200 µl/min with the loading buffer (0.1 % formic acid). Peptides were trapped and desalted on-line on a peptide microtrap (Phenomenex UHPLC Fully porous polar C18, 2.1 mm) for 3 min before elution onto an analytical column (Phenomenex Luna Omega Polar C18 analytical column 1.6 µm, 100 x 1.0 mm, 100 Å), maintained at 2°C. A 6-min linear-gradient (10%-45%) with 80% acetonitrile with 0.1% formic acid was used to separate the peptides (Agilent 1290). The eluate from the analytical column was directed into the TOF mass spectrometer (Bruker) with electrospray ionization. Blank injections were performed between each sample injection to prevent the carryover of peptides between runs. HDX samples were analyzed in MS mode without trapped ion mobility.

Tandem mass spectra were searched using MASCOT against the cRAP and contaminants protein database (http://ftp.thegpm.org/fasta/cRAP), containing the sequences of ApoE3 and ApoE4, with precursor ion mass tolerance set at 10 ppm and fragment ion mass tolerance at 0.05 Da. No enzyme specificity was applied, with a maximum allowance of two missed cleavages, and no fixed or variable modifications. The false discovery rate at the peptide identification level was set to 1%. The peptide map was prepared using the DigDig software, version 0.8.1^28^ and peptide redundancy was calculated. The MS raw files of deuterated and undeuterated samples, along with the list of peptides and their parameters (score, retention time, charge, sequence, and mobility), were analyzed using HDExaminer version 3.4.1 (Sierra Analytics, Modesto, CA).^29^ The software analyzed peptide-level deuteration over time and calculated the score based on the fit of theoretical and actual isotope clusters. Confidence levels were assigned as high, medium, or low. Peptides were manually reviewed to improve the fit, and only high- and medium-confidence peptides, up to 20 amino acids in length, were included in further calculations.

PyHDX v0.4.3^30^ was then used to calculate relative fractional uptake (RFU), Gibbs free energy (ΔG), and differential Gibbs free energy ΔΔG values, allowing comparison of protein exchange kinetics across states. RFU values for free ApoE3/4 and ligand-bound forms were calculated by weighted averaging, where each residue’s RFU was determined by averaging RFUs of overlapping peptides, weighted by the inverse peptide length, and visualized in a linear bar plot. Residue-level ΔG values were derived using the Linderstrøm-Lang model, which describes amide hydrogens transitioning between exchange-incompetent “closed” and exchange-competent “open” states. ΔG values were calculated using HDX-MS exchange rates (k_obs) under specified parameters: temperature (275.15 K), pH (7.1), stop loss (0.000005), stop patience (50), learning rate (20000), momentum (0.5), epochs (300,000), with regularizers set to 1 and 2.5, respectively.

### Stem cell culture

Two isogenic human iPSC lines used in this study were passaged and maintained using standard feeder-free culture protocols. In brief, feeder-free cultures were grown on Matrigel-coated plates (Corning) in mTeSR^TM^1 medium (STEMCELL Technologies) supplemented with half of the recommended dose of ZellShield® (Minerva Biolabs). Cells were passaged using 0.5 mM EDTA (Thermo Fisher Scientific) in PBS or manually. The “E4” cell line was derived from a patient with a sporadic form of AD with ApoE4/4 status. The isogenic cell line “E3” was obtained by correction of ApoE4/4 to ApoE3/3. Both isogenic iPSC lines were kindly provided by Dr. Li-Huei Tsai and described previously.^31^

### CO culture and TRN treatment

COs were generated using the protocol described previously.^32,33^ Briefly, for spheroid formation, cells were plated at day 0 into non-adherent V-shaped 96-well plates at 2000– 3000 cells in 150 µl of mTeSR^TM^1 medium with 50 µM ROCK inhibitor (S1049, Selleckchem). Plates were centrifuged for 2 min at 200 *g* to facilitate spheroid formation. Non-adherent cell culture plates were prepared with poly(2-hydroxyethyl methacrylate) (poly-HEMA; P3932, Merck) coating. On day 2, the cell culture medium was exchanged for fresh mTeSR^TM^1 without ROCK inhibitor. When spheroids reached the size of 400– 600 μm, fresh Neural Induction Medium was added every day for six days (usually from day 3 to day 8).^34^ The next day, twelve organoids were transferred to one 6 cm cell culture dish. The remaining medium was aspirated, and dry organoids were embedded in 7 µl of cold Geltrex™ (Thermo Fisher). Geltrex™ was left to solidify as hanging drops on the inverted cell culture dish for 10 min at 37 °C. Solidified Geltrex™ drops with organoids were gently detached from the bottom and cultured without shaking in Cerebral Organoid Differentiation Medium (CODM)^34^ without vitamin A for seven days. Subsequently, organoids were cultured in CODM with vitamin A and were moved on an orbital shaker on day 26 (± two days). CODM was changed three times a week by aspirating at least half of the old medium and replacing it with a fresh medium. ZellShield® (Minerva Biolabs) was used to prevent contamination in all media. The treatment with TRN or TMP was performed for 50 days, from day 50 (D50) to day 100 (D100). During the treatment, CODM was continuously supplemented with 100 µM TRN or 100 µM tramiprosate diluted in the cell culture medium. Samples were harvested from two independent batches of CO differentiation. Control samples (NTR) were treated with medium only.

### Cryo-sections, immunohistochemistry (IHC) and microscopic analysis

Harvested COs were fixed with 3.7% paraformaldehyde for 1 h and washed with PBS. For cryo-sections, fixed organoids were saturated with 30% sucrose (Merck), embedded in O.C.T. medium (Tissue-Tek), and frozen. Then, 10 µm sections were prepared on cryostat Leica 1850. Excessing O.C.T. medium was removed by 15 min PBS wash prior to IHC staining. Sections were permeabilized in 0.2% Triton-X (Merck) in PBS and blocked in 2% normal goat serum (Merck) in permeabilization solution. Samples were incubated with primary antibodies in a blocking solution at 4 °C overnight followed by incubation with secondary antibodies (Alexa FluorTM; Thremo Fischer Scientific) for 1 h at room temperature. Nuclei were visualized by Hoechst 33342 (Thermo Fischer Scientific). The following primary antibodies were used: NFL (# 2837; CST), MAP2 (#8707; CST), SYN1 (#5297; CST), GFAP (#12389; CST), S100β (ab11178; Abcam), and IBA1 (#019-19741; Labmark). Samples were imaged with the inverted microscope Zeiss Axio Observer.Z1 with confocal unit LSM 800.

### Sample preparation for proteomic assays

For proteomic assays, individual organoids were washed with PBS on D100, incubated in Cell Recovery Solution (Corning) for 1 h, washed with PBS, and stored at -80 °C. Five to six organoids from three individual batches were harvested for analyses.

### RNA isolation and mRNA sequencing

Harvested organoids were washed with PBS, lysed with 1 ml RNA Blue reagent (Top-Bio), and stored at -80 °C. Total RNA was isolated from three biological replicates (ApoE3/3=E3; ApoE4/4=E4) for each condition (NTR, TMP, TRN) with Direct-zol RNA Microprep kit (ZymoResearch) according to the manufacturer’s instructions. RNA quality was assessed by TapeStation 2200 (Agilent Technologies; RNA Screen Tape), and only samples with RINe values ≥ 8.5 were used for library preparation. Poly-A selected libraries were made from 300 ng of total RNA using QuantSeq FWD 3’mRNA Library Prep Kit (Lexogen) in combination with UMI Second Strand Synthesis Module for QuantSeq FWD and Lexogen i5 6 nt Unique Dual Indexing Add-on Kit (Lexogen) with 14 - 18x PCR cycles according to the manufacturer’s instructions. Quality control of library quantity and size distribution was done using QuantiFluor dsDNA System (Promega) and High Sensitivity NGS Fragment Analysis Kit (Agilent Technologies). The final library pool was sequenced on NextSeq 500 (Illumina) using High Output Kit v2.5 75 Cycles in single-end mode, resulting in an average of 10 million reads per sample.

### Alignment of RNA-seq samples and quality control

The raw reads were initially trimmed using Trimmomatic^35^ and subsequently mapped to the human reference genome hg38 with STAR (v.2.7.3).^36^ Transcript-level expression quantification was carried out using RSEM,^36^ with results aggregated at the gene level according to GENCODE (v.30).^37^ RNA-SeqQC^38^ and *Picard* (v2.2.4) tools were employed to assess the quality control metrics, which included the number of aligned reads (minimum average of 8.740 ± 0.593 million), the fractions of duplicate reads (0.723 ± 0.057), the fractions of uniquely mapped reads (0.75 ± 0.020), and the mean GC content (40.597% ± 0.630%). No systematic differences were observed among the replicates or across the samples.

### Differential gene expression analysis

To identify genes that showed differential expression between treatments (NTR/TRN/TMP), a differential gene expression (DGE) analysis was performed. A count matrix was constructed jointly for all samples to include genes where counts per million (CPM) were ≧ 4 in at least 15% of the samples. The read count matrix was adjusted for technical confounders, i.e., sequencing batch and two principal components (PC1 and PC2) calculated on the matrix of 15 sample-level covariates including the fractions of intronic/intergenic/mRNA/coding/ribosomal bases, fractions of uniquely mapped reads / duplicate reads, mean GC content, estimated fractions of GABAergic neurons / glutamatergic neurons / astrocytes / oligodendrocytes, the fractions of uniquely mapped reads, chimeric reads and too-short reads. Differential analysis was performed using the *dream* function from the variancePartition R package (v1.0.27).^39,40^

### Visualisation and further analysis of mRNA sequencing data

To further visualise the effect of TRN on COs, differential gene expression data processed as described previously were filtered and genes with p-adj<0.1, and absolute log fold change abs(log2FCH)>0.6 in between given conditions excluding novel transcripts were used for further analyses (n=206). Genes changed in ApoE4 in any condition (E4_NTR, E4_TRN, E4_TMP) compared to ApoE3 (E3_NTR) were plotted and tested based on the normality of the data with appropriate statistical tests using GraphPad Prism v8 (GraphPad Software). The biological processes influenced by the changes in gene expression were analysed using g:profileR^41^ (g:Profiler version e111_eg58_p18_b51d8f08) as described previously,^42^.

### Liquid chromatography-tandem mass spectrometry protein analysis (LC-MS/MS)

CO tissue samples were processed using the protocol and analyzed using the LC-MS/MS assay (including data processing) as described previously.^5^ The following reagents were used: LC/MS grade Acetonitrile (ACN, 34967), Isopropanol (IPA, 34965), and Formic acid (FA) were obtained from Honeywell (Charlotte, USA), Ammonium bicarbonate (AmBic, BioUltra, ≥ 99.5% purity, 09830), Sodium deoxycholate (SDC, BioXtra, ≥ 98.0% purity, 30970), Iodoacetamide (IAA, ≥ 99% purity, I6125) from Sigma Aldrich (St. Louis, MO, USA), 1,4-dithiothreitol (DTT, ≥ 99% purity, 6908.1) from Carl Roth GmbH + Co. KG (Karlsruhe, Germany), Trypsin gold, Mass Spec Grade, from Promega (Madison, WI, USA), Pierce BCA Protein Assay Kit reagents from ThermoFisher Scientific (Waltham, MA, USA). Synthetic isotopically labeled (SIL) peptide standards (SpikeTides_L crude) were synthesized by JPT Peptide Technologies Inc. (Acton, MA, USA). The ultrapure water was prepared using the water purification system (arium® Comfort System, Sartorius).

COs (stored at -80 °C) were lyophilized using SpeedVac vacuum concentrator (Savant SDP121 P, ThermoFisher Scientific, USA) and homogenized with glass bead (BeadBlasterTM 24, Benchmark Scientific, Edison, NJ, USA). Proteins were precipitated with 80% IPA (100 µl). Protein pellets formed after vortexing (VELP Scientifica), sonication (37 Hz, 5 min; Elmasonic P, Elma Schmidbauer GmbH) and mixing (200 rpm, 10 min; HeidolphTM MultiReax) was centrifuged (12.3 RCF, 5 min; Micro-Star 12, VWR®, Radnor, PA, USA) and dried in SpeedVack after removal of the supernatant. The dried protein pellets were solubilized by adding 100 µl AmBic buffer (100 mM) with SDC (5 mg/ml) (AmBic+SDC). All samples were vortexed (10 s, 2000 rpm), homogenized (4 m/s, 10 s, two cycles with 10 s inter-time), sonicated (1 min, 80 kHz, Elmasonic P, Elma Schmidbauer GmbH), mixed (10 min, 2035 rpm) and centrifuged (1 min, 12,300 RCF). The total protein content (µg/ml) in each protein homogenate sample was measured using the BCA protein assay. The protein concentration in each sample was adjusted to 0.1–0.3 µg/µl by adding the AmBiC+SDC buffer. After centrifugation, aliquots of 60 µl (6–18 µg of total protein) were reduced with 20mM DTT in 50 mM AmBic (10 min; 95 °C) and alkylated with 40mM IAA in 50 mM AmBic (30 min; room temperature in the dark). The remaining homogenates from individual COs were mixed to prepare a quality control (QC) sample for calibration curve measurements. The samples were digested with trypsin (1:20–40, enzyme: total protein, w/w), sealed and incubated (37 °C; 16 h) with gentle shaking. The digestion was stopped by acidification with 2% FA (200 µl), and SIL peptide standards were added to the samples in a final concentration of ≈22–29 nmol/l. Peptide mixtures were then purified using the solid-phase extraction (SPE) method (mixed-mode cartridge, Oasis® PRiME HLB – 30 mg, Waters Corp. Milford, MA, USA): after loading on the column sorbent, the samples were washed with 2% FA, eluted with 500 µl of 50% ACN with 2% FA and dried in SpeedVac.

The samples were reconstituted in 5% ACN with 0.1% FA (15–20 µl). Analysis of peptides was performed on the LC-MS/MS system (UHPLC Agilent 1290 Infinity II coupled to a triple quadrupole mass spectrometer Agilent 6495B, Agilent Technologies, CA, USA) operated in a positive ion detection mode. Samples (3 µl) were injected into the C18 analytical column (Acquity UPLC CSH 1.7 μm, 2.1 mm x 100 mm, Waters Corporation, MA, USA). The gradient mode of mobile phases A (0.1% FA) and B (0.1% FA in 95% ACN) was: initial 5% B; 25 min 30% B; 25.5 min 95% B; 30 min 95% B; and from 31 to 35 min with 5% B; flow rate was set to 0.3 ml/min. The electrospray ionization (ESI) source parameters were: temperature = 200 °C, capillary voltage = 3500 V. The same multiplex selected reaction monitoring (SRM) assay was used for the detection and relative quantification of protein targets in the COs as published in the previous study.^5^ Manually inspected data were processed in Skyline software (version 21.2.0.369, MacCoss Lab, UW, USA). Peptide transitions with a reproducible signal (on average, %CV < 15%) selected for the relative quantification of proteins shown in the study are listed in **Table S1**. Calculated protein concentrations using the peak area of the spiked SIL peptide standards were normalized to the GAPDH levels. The final results were visualized as fold-change values in this study, i.e. analyzed protein levels in TRN-treated E3 and E4 organoids (E3_TRN, E4_TRN) were related to the corresponding non-treated (NTR) E3 and E4 organoids (E3_NTR, E4_NTR) in each batch to see the effect of TRN protein expression relative to protein levels found in organoids without the treatment.

## Results

### TRN promotes ApoE4 T-shaped dimer dissociation through electrostatic interactions

To assess the TRN-ApoE4 interaction, we simulated the dissociation of ApoE dimers, using adaptive sampling molecular dynamics (MD) simulations, starting from a T-shaped dimeric state of ApoE3 and ApoE4 (**Figure 2A**), alone or in the presence of 100 small molecules (TRN or SPA).

**Figure 2.**
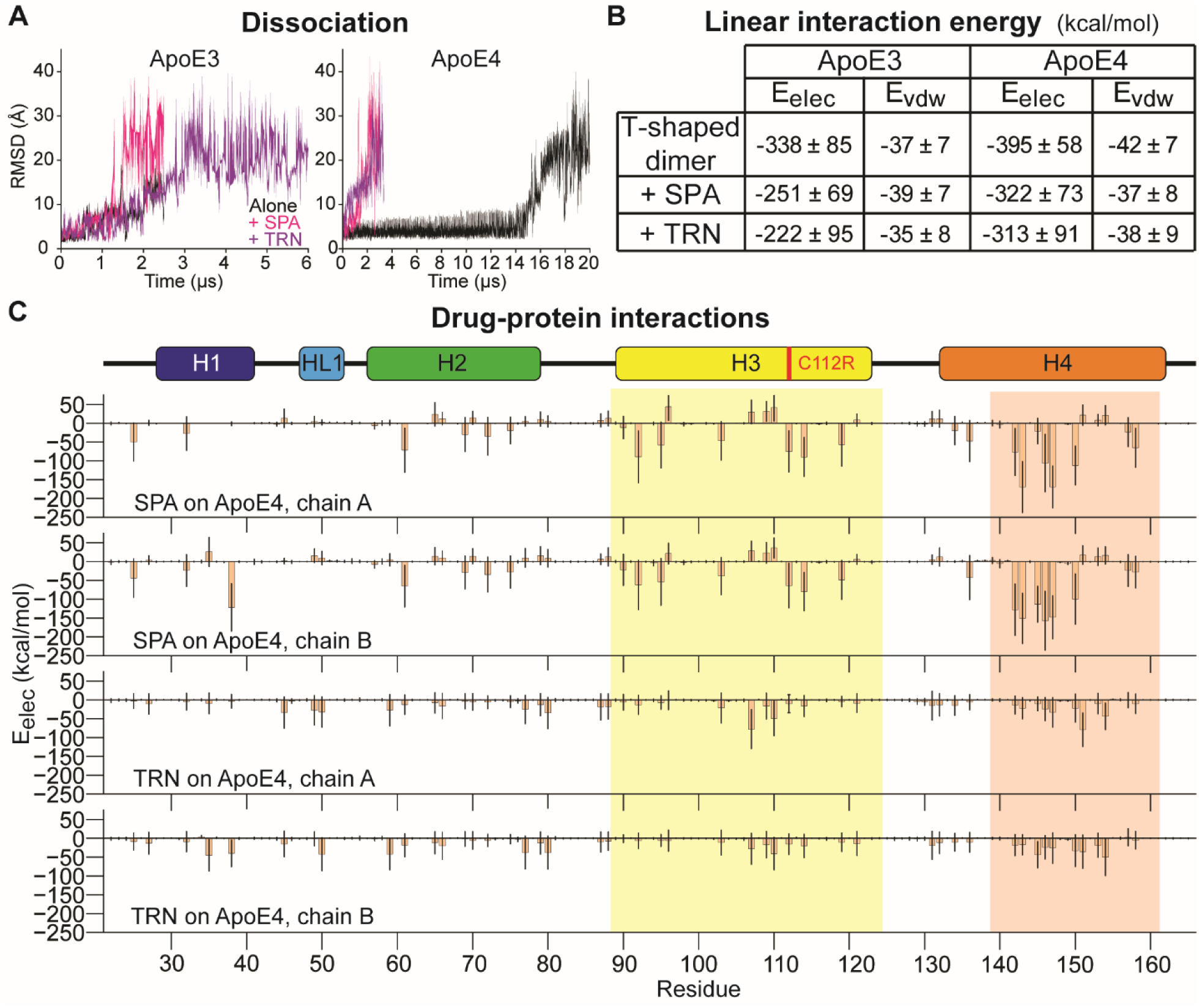
TRN and SPA bind to ApoE T-shaped dimers and promote their dissociation. (A) RMSD of Cɑ atoms during the dissociation adaptive MD simulations of ApoE3 (left) and ApoE4 (right) T-shaped dimers, alone (black) or in the presence of TRN (purple) or SPA (pink). (B) Electrostatic (E_elec_) and van der Waals (E_vdw_) components of the linear interaction energy (LIE, in kcal/mol) between the two protomer chains of ApoE3 and ApoE4 T-shaped dimers, alone or in the presence of SPA or TRN molecules, during the adaptive MDs; the uncertainty values represent the standard deviation of the mean energies in the initial 500 ns (5000 snapshots). (C) Electrostatic component (E_elec_) of LIE between TRN or SPA with every residue of each chain (A and B, as defined in Figure 1B) of ApoE4 T-shaped dimer, during the adaptive MDs; the uncertainty bars represent the standard deviation of the mean energies during the initial 500 ns (5000 snapshots). Colored boxes highlight regions where TRN and SPA preferentially bind. See **Supplementary Figure S2** for data with ApoE3.

We measured the evolution of the root-mean-square deviation (RMSD) of the protein Cα atoms with respect to their starting structures, setting the threshold for tracking the dimer dissociation to RMSD values above 15 Å. All systems (ApoE4 with SPA or TRN, free ApoE3) showed dissociation after ca. 1.5 – 3 μs of combined simulation times (**Figure 2A**), while the free ApoE4 took one order of magnitude longer to dissociate (16 µs). These results suggest that both TRN and SPA can exert a similar effect in promoting the dissociation of ApoE4 T-shaped dimers.

The interface interactions in the ApoE T-shaped dimers, as assessed by the linear interaction energies, LIE^26,43^ during the initial part of the adaptive simulations (first 500 ns), are mainly driven by electrostatic interactions, which were nearly one order of magnitude stronger than the van der Waals component (**Figure 2B**). These energies were stronger for ApoE4 than for ApoE3, which agrees with the observation that the dimer dissociation was much slower for ApoE4 than for ApoE3. The presence of TRN or SPA significantly decreased the electrostatic interactions (for ApoE4, by ca. 82 kcal/mol and 73 kcal/mol, respectively), which may explain the higher dissociation propensity of the dimers in the presence of those small molecules. To understand the effects of TRN and SPA at the molecular level, we assessed the binding energies of those small molecules to the amino acids of dimeric ApoE3 and ApoE4, during the same simulations. The van der Waals component of LIE was negligible compared to the electrostatic component. The results also revealed that TRN binds to the same regions of the dimers as SPA but with weaker interaction energies (**Figure 2C and S1**). Most of the dimer-interface residues in chain A are positively charged, while those in chain B are negatively charged. Total interaction energies were stronger with SPA (more negative) than with TRN, although TRN interacted with more residues than SPA. This is because TRN has one positive and one negative moiety, while SPA has two negative moieties. The variability of interaction energies is large for both TRN and SPA due to the nonspecific nature of those interactions, which results in a relatively fast on/off switch of the contacts between the ligands and the residues. The strong electrostatic interactions of TRN and SPA with the charged dimer-interface residues interfered with the protein-protein interactions in that region (**Figure 2C**), weakening them and ultimately contributing to their faster dissociation. These effects have previously been observed for SPA.^5^

Next, we tested the effect of SPA and TRN on ApoE dimer association with adaptive goal MD simulations. Monomeric frames were defined as those in which the minimum distance between heavy atoms of different chains is at least 8 Å. This observation was robust to the selection of the distance threshold (**Figure 3A, Figure S2**). The proportion of monomeric frames increased by a factor of ca. 2 in the presence of either SPA or TRN, indicating that both molecules can suppress the association of ApoE dimers. VAMP classification of dimeric states into six clusters revealed a plethora of different ApoE dimeric conformations, including parallel, V-shaped, T-shaped, and anti-T-shaped dimers (**Figures 3A-B, Table S2, Figure S3**).

**Figure 3.**
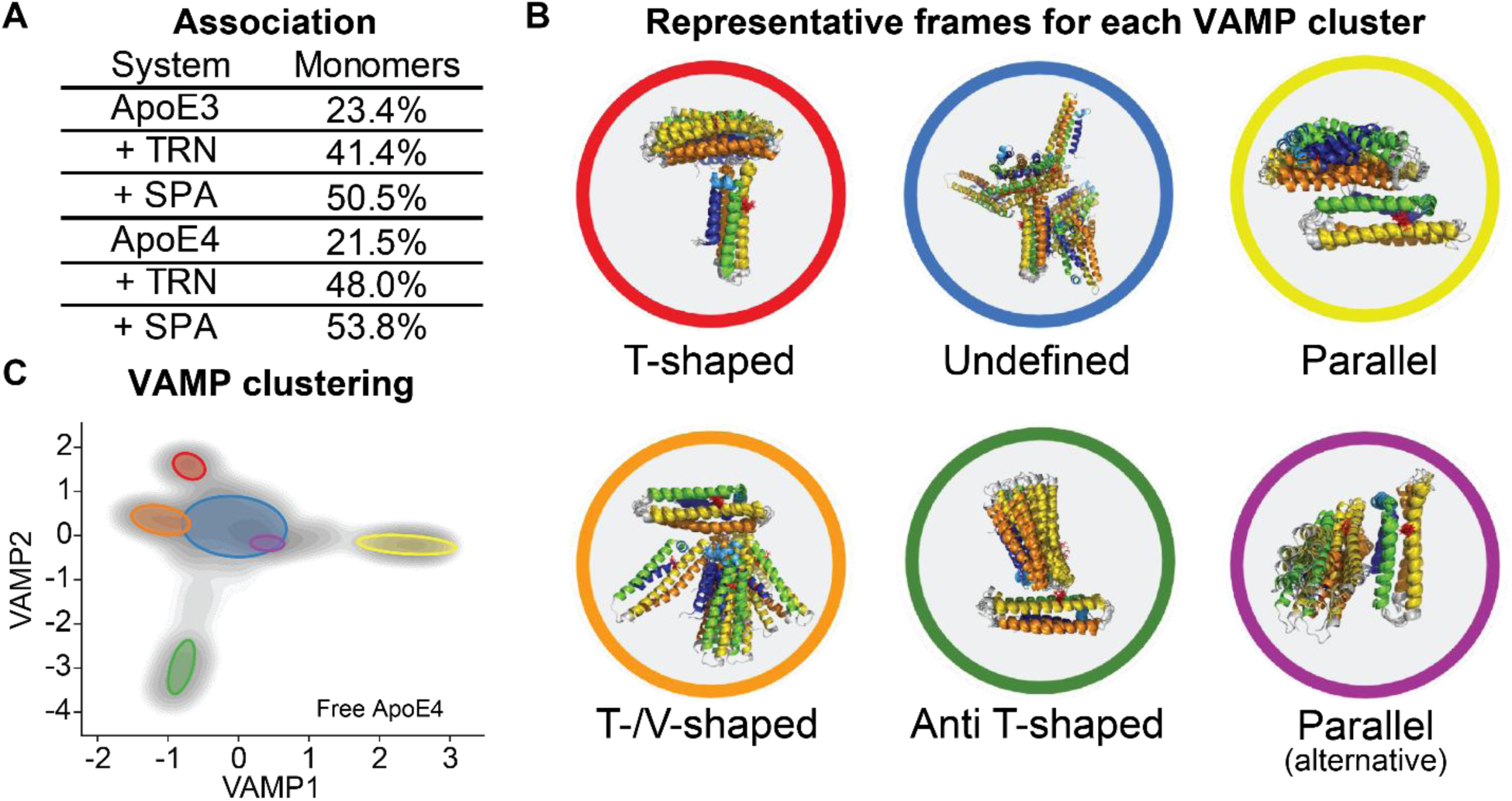
VAMP classification of ApoE dimers and inhibition by TRN and SPA. (D) Percentage of monomeric states in all systems. (E) Representative frames and dimer type for each VAMP cluster of free ApoE4 (as described in (F)). (F) Free energy landscape (grey shades) and VAMP clusters (represented by the coloured ellipses) of the dimeric states of free ApoE4 during the association adaptive goal MDs. Structures are colored as described in Figure 1.

We present the VAMP clustering results only for the most aggregation-prone system, i.e. the free ApoE4 system, which we used to determine the conformations of interest. To the best of our knowledge, only the T-shaped, V-shaped and parallel dimers have been observed in crystallographic structures.^5^ Previously, we experimentally identified T- and V-shaped dimers as protein-protein crystallographic dimers, then later confirmed as native protein dimers by mutagenesis. The difference between these two dimers lies in the angle between the two chains. Likewise, parallel crystallographic dimers were also observed in the simulations but have not been validated experimentally.

The VAMP analysis identified the parallel dimer as forming the largest cluster observed in the free ApoE4 system, consisting of 11.5% of all frames in the adaptive goal simulation (**Figure 3A-B**). The RMSD analysis confirmed that 7.6% of the free ApoE4 frames had RMSD values below 6 Å when compared to the crystallographic parallel dimer. This percentage decreased to 1.7% in the presence of TRN and dropped to 0% with SPA, suggesting a higher efficacy of SPA at preventing parallel dimer formation. Both SPA and TRN formed favorable interactions with helix H4 (**Figure 2C**), which is also a critical component of the parallel dimer interface (**Figure S4**). Interestingly, only small or inexistent populations of parallel dimers were observed in the ApoE3 systems (0.5% for free ApoE3, 0.0% for ApoE3 with TRN, and 0.1% for ApoE3 with SPA) (**Table S2**). We observed the formation of a dimer, which we termed “anti-T-shaped dimer” concerning the T-shaped dimer previously described,^5^ since chain B interacts with the opposite side of chain A (**Figure 3B**). A key observation is that the mutated residue at position 112 (C112 for ApoE3, R112 for ApoE4) appears to be involved in the interface of this dimer. Notably, an anti-T-shaped VAMP cluster was detected only in the free ApoE4 system (**Figure 3B**), and the RMSD analysis confirmed that these interactions were unique to this system (**Table S2**). While interactions involving the mutated residue at position 112 could be of particular interest, potentially offering direct insights into the mutational effects from ApoE3 to ApoE4, this anti-T-shaped dimer has never been observed experimentally thus far.

Altogether, our simulations demonstrated that SPA and TRN effectively increase the population of monomeric species, indicating their ability to prevent ApoE dimer association. VAMP clustering revealed diverse ApoE dimer conformations, with the fractions of the parallel dimer of ApoE4 being significantly reduced in the presence of SPA and TRN. Both molecules interact favorably with helix H4, crucial for parallel dimer formation. Additionally, a unique anti-T-shaped dimer was identified in the ApoE4 system, involving the mutated residue at position 112, although this finding remains to be validated experimentally.

### TRN suppresses ApoE4 aggregation and modifies its dynamics

While molecular dynamics provides insights into the initial stages of the ApoE aggregation, the simulation of the entire aggregation process involving dozens or possibly hundreds of protomers is computationally too demanding. Therefore, we utilized static light scattering (SLS) to experimentally monitor protein aggregation in real-time.

Full-length ApoE3 or ApoE4 (10 µM) was mixed with a 2000-fold excess of SPA, a metabolite of the prodrug TMP,^5^ or with TRN. As expected, ApoE3 showed minimal aggregation, whereas ApoE4 aggregated rapidly. TRN and SPA both significantly suppressed ApoE4 aggregation (**Figure 4A**). To quantify the extent of ApoE aggregation inhibition, we fitted the aggregation curves analytically with a single exponential function with a linear drift component **(Figure 4B, Table S3)**. The amplitude of the initial exponential aggregation phase (*A*_1_), which describes the amount of aggregated ApoE in the initial exponential aggregation phase, was significantly reduced in the presence of SPA (Δ = 73 ± 10 % less aggregation) or TRN (Δ = 40 ± 7 % less aggregation), as compared to ApoE4 alone (p-values < 0.0001). This result confirms that both compounds can decrease ApoE4 aggregation, with SPA having a stronger inhibitory effect.

**Figure 4.**
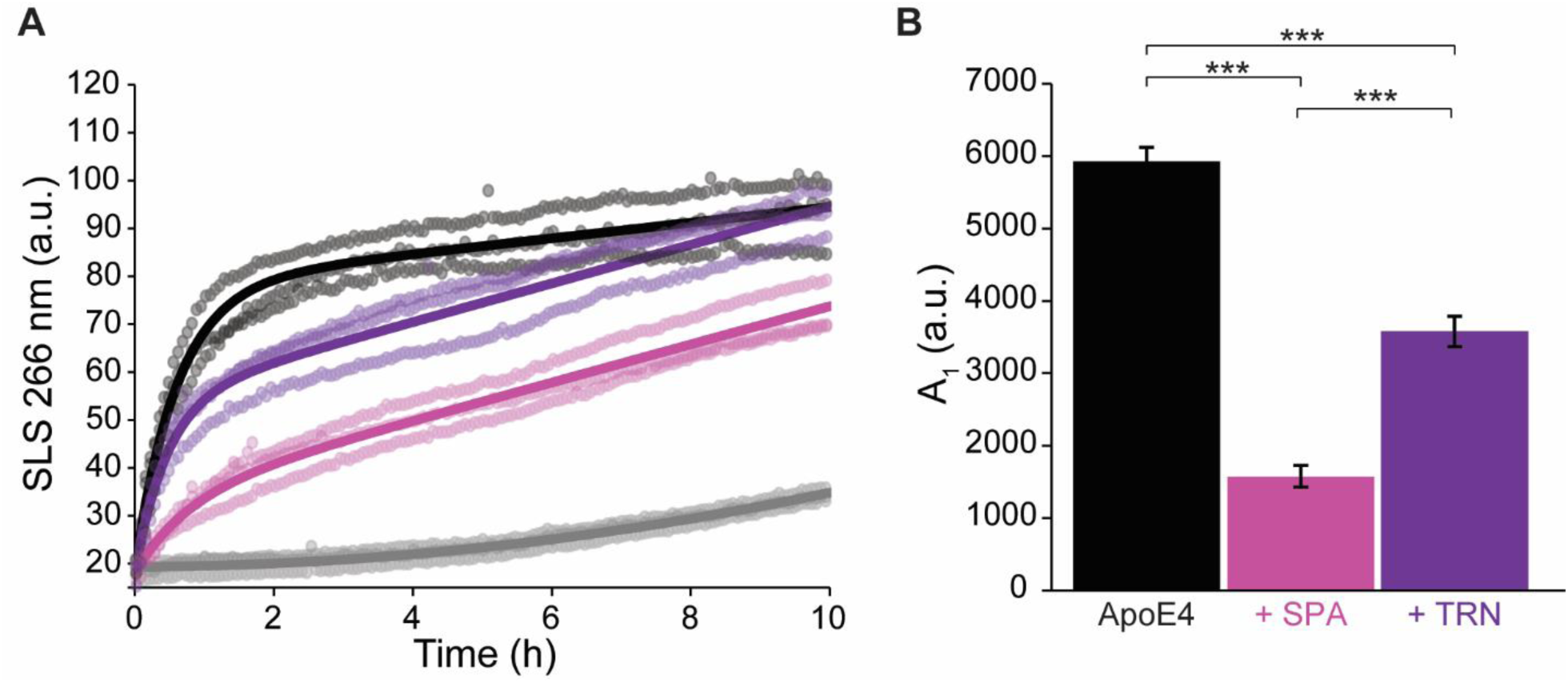
TRN and SPA suppress ApoE4 aggregation and solvation. (A) Static light scattering (SLS) of ApoE3 (grey), free ApoE4 (black) and ApoE4 with a 2000-fold excess of SPA (pink) or TRN (purple). The data was collected in 10 mM Tris 50 mM NaCl pH =7.4 buffer at 37°C. (B) Fitted amplitudes of initial aggregation phase (*A*_1_) using a single exponential function with linear drift component (*Eq. 1*). All fitted parameters are given in **Table S1**. Error bars represent standard errors. ***: p-value < 0.0001 for T-test.

To further evaluate the effect of TRN on protein dynamics, dimerization, and aggregation, we conducted hydrogen-deuterium exchange coupled with mass spectrometry (HDX-MS) experiments. The peptide mapping confirmed complete sequence coverage, demonstrating 100% coverage of the protein sequences with 840 unique peptides for ApoE3 and 857 unique peptides for ApoE4. The peptide coverage map and redundancy over the sequence are illustrated in **Figures S5-7**.

At 60 s, SPA and TRN reduced the RFU across the entire ApoE4 sequence, with a similar trend observed at later time points, as depicted by the RFU over sequence and the heat map **(Figure 5A, Figure S8-9)**. ΔΔG calculations (**Figure 5B-D**) reveal that in the presence of TRN, ApoE4 showed significant changes in regions 8-21 (N-terminal), 72-76 (H2 residues), 87-97 (N-terminal of H3), 114-115 (near the mutation site in H3), 128-132 (N-terminal of H4), 162-235, and 284-300 (C-terminal membrane-binding domain). A similar trend of ΔΔG was observed for ApoE4 with SPA, except for the residues 72-76 (H2). In contrast, no ΔΔG changes were detected in ApoE3 with SPA or TRN, suggesting that TRN and SPA specifically induce a localized reduction in ApoE4 flexibility (**Figure S10**). The mean square error loss for ΔΔG calculations is included in the supplementary material (**Figure S11**).

**Figure 5.**
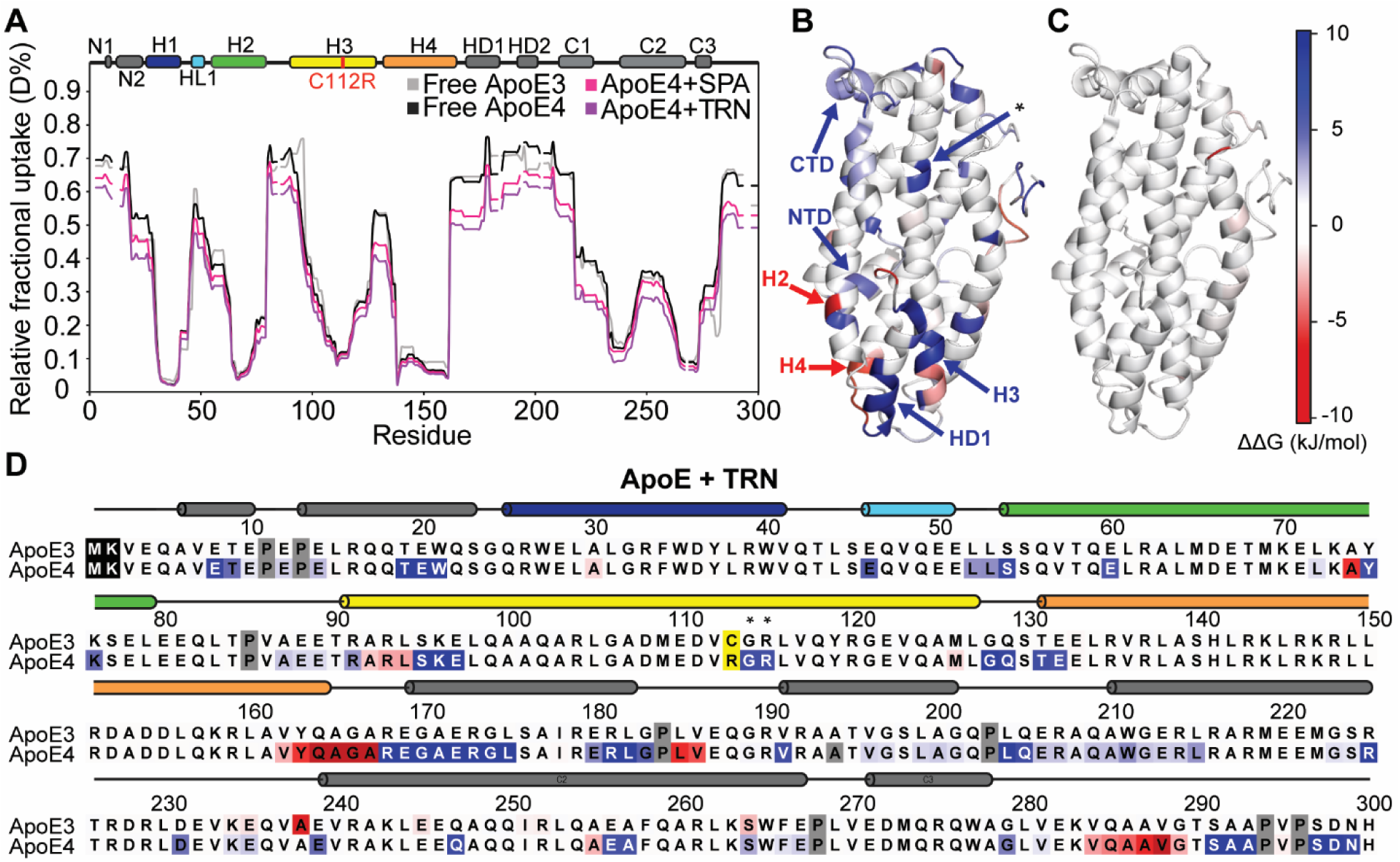
HDX-MS of ApoE3 and ApoE4 with SPA and TRN. (A) Relative fractional uptake (RFU) over amino acid residues for ApoE3 free, ApoE4 free, ApoE4 with SPA, and ApoE4 with TRN after 60 s of deuterium exposure represented in grey, black, pink, and purple, respectively (see legend). Data were collected at four time intervals: 60 s, 120 s, 600 s, and 1800 s. (B) ΔΔG values of ApoE4 with TRN, and (C) ApoE3 with TRN, mapped on the structure of ApoE (PDB 2L7B), showing regions of altered stability. (D) ΔΔG values mapped on the amino acid sequences of ApoE3 and ApoE4, with values compared to the free ApoE3/4 states. Differences are highlighted in red and blue. Red regions indicate lower ΔΔG values, suggesting increased amide hydrogen (H) to deuterium (D) exchange compared to the free protein, while blue regions indicate reduced exchange. The mutation site (C112R) is highlighted in yellow with an asterisk, and the first two residues, excluded from pyHDX analysis, are shown in black. Proline residues, non-exchangeable in HDX-MS, are indicated in grey.

### TRN modifies the expression profile of ApoE4/E4 in organoids

To explore the effect of TRN on a human-relevant cellular model, we used induced pluripotent stem cell (iPSC)-derived COs differentiated from an isogenic pair of cell lines carrying *ApoE3/E3* or *ApoE4/E4* genotype.^31^ As schematized in **Figure 6A**, both cell lines were used to generate mature COs with an organized morphology and expressing standard neuronal and glial markers, i.e., NFL, MAP2, SYN1, and S100b, GFAP, respectively, including a marker of microglia, IBA1 (**Figure 6B**). At day 50 of *in vitro* culture, 100 µM TRN or TMP, a precursor of SPA,^5^ was added to the cell culture media of both *ApoE3/E3* and *ApoE4/E4* organoids and maintained in the organoid culture media for another 50 days, with media exchange every 2-3 days. On day 100 of *in vitro* culture, organoids were harvested for transcriptomic and proteomic analyses. All experiments were performed in three independent batches, and data were processed as described in the methods.

**Figure 6.**
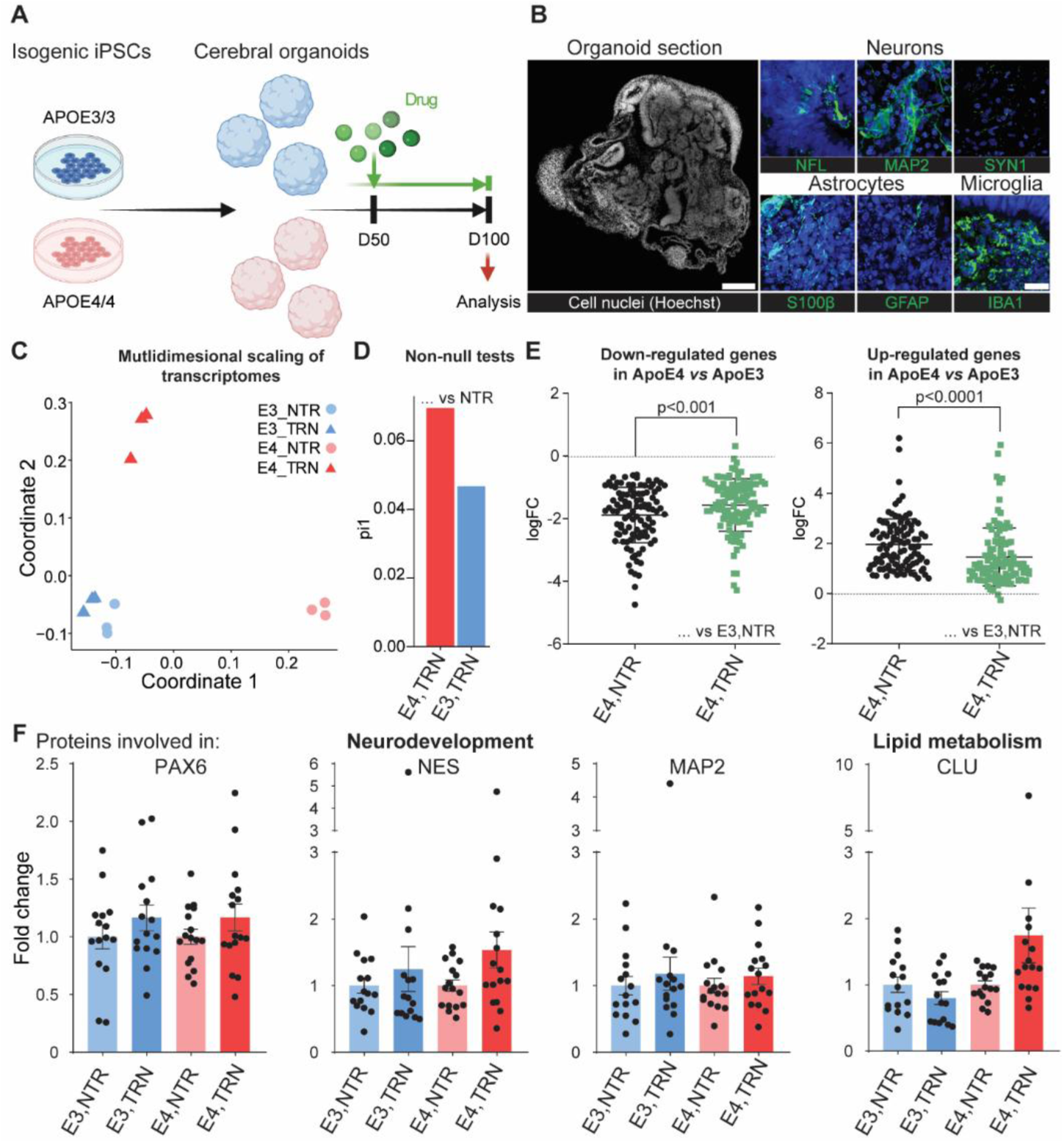
Effects of TRN treatment on COs. (A) Schematic representation of experimental design. (B) Paraffin sections from D100 organoids show overall morphology and representative staining of neuronal and glial markers. Left scale bar: 500 µm, right scale bar: 50 µm. (C-D) Characterisation of transcriptomic data based on next-generation sequencing screening (mRNA sequencing) of non-treated control COs (NTR) and COs treated with TRN. (C) Multidimensional scaling. (D) Proportions of non-null tests (pi1) for various scenarios. Higher estimates indicate more significant differences. (E) LogFoldChange expression of genes differentially expressed in between *ApoE4/ApoE3* genotypes (p-adj<0.1, abs(log2FCH)>0.6; n=206) and their shift in expression after TRN treatment. *Left:* down-regulated genes in ApoE4 compared to ApoE3 and shift of their expression toward the ApoE3 after the TRN treatment (p<0.001; n=104). *Right:* up-regulated genes in ApoE4 compared to ApoE3 and shift of their expression toward the ApoE3 after the TRN treatment (p<0.0001; n=102). (F) Changes in protein levels as a function of genotype and TRN treatment. NTR *ApoE3/ApoE3* COs protein level is defined as 1. Protein levels in the ApoE3 and ApoE4 TRN-treated COs is relative to protein levels expressed in respective non-treated (NTR) ApoE3 and Apoe4 COs. Each dot represents one organoid, n=5-6 organoids from three independent batches were analyzed and are visualized here.

Interestingly, our data analysis revealed that TRN treatment specifically affects the biology of *ApoE4/E4* organoids while having only a minor impact on *ApoE3/E3* organoids. This observation is in excellent agreement with the *in vitro* HDX-MS experiments (**Figure 3A**). As shown in **Figure 6C**, the transcriptomic data, adjusted for technical confounders (**Figure S12A**), was used for clustering and resulted in a distinct separation between *ApoE3/E3* and *ApoE4/E4* COs (**Figure 6C**). Treatment with TRN only had a significant effect on monozygotic COs bearing the ApoE4 allele, which is the first genetic risk factor for late-onset AD.^2^ This was further validated by a higher proportion of non-null tests for treatment comparisons within *ApoE4/E4* COs compared to ApoE3/E3 COs (**Figure 6D; Table S3**). Interestingly, although TRN did not explicitly regulate a specific pathway, it induced a notable shift in the expression of over 200 genes, bringing gene expression of *ApoE4/E4* COs closer to that of ApoE3/E3 COs (p<0.001) (**Figure 6E, Figures S11C-D, Tables S4-6**).

A similar expression trend was observed with TMP treatment,^5^ acting here as a positive control, with several of the identified genes associated with neurodevelopment, autophagy pathways, or lipid metabolism (**Figure S13, Table S6**). Importantly, this upregulation trend was also confirmed on a protein level by mass spectrometry (**Figure 6F**), as shown by proteins related to neuronal development (PAX6, NES, MAP2) and lipid metabolism (CLU). Collectively, our data suggest that TRN can significantly modulate gene expression in *ApoE4/E4* organoids, aligning it more closely with the gene expression profile of *ApoE3/E3*, particularly in genes linked to neurodevelopment, lipid metabolism, and autophagy, all pathways previously connected to AD pathogenesis.

## Discussion

The “ApoE Cascade Hypothesis” stipulates that the aggregation of ApoE would play an important role in neurodegeneration.^2^ Indeed, different ApoE isoforms, as well as defects in ApoE oligomerization and lipidation states, are involved in many ApoE-related neurodegenerative pathologies, such as AD, PD, LBD, and macular degeneration. ApoE is mostly lipidated^44^, and a lack of lipidation is correlated with increased Aβ aggregation *in vivo*. ApoE4 is the least-lipidated, the most aggregation-prone isoform, and a genetic risk factor for early-onset AD, PD, and LBD. Conversely, ApoE3 and ApoE2 are less aggregation-prone and more lipidated^2,4^. In the case of AD, ApoE binds to toxic Aβ fibrils, and inhibits their elongation.^45^ A notable exception occurs in the context of macular degeneration, where ApoE2 is a genetic risk factor, and ApoE4 is a genetic protective factor.^3^

ApoE3 and ApoE4 only differ by a single mutation, C112R. The effect of this mutation is profound in the ApoE4 T-shaped dimers, in which the R61-E109 interaction is lost, and the Q123 changes conformation, which leads to the destabilization and bending of helix 3.^5^ In our previous study, we showed that TMP could directly inhibit ApoE4 aggregation *in vitro* using purified proteins and strongly affect lipid compositions *in vivo* in COs. However, TMP is metabolized *in vivo* into SPA, suppressing ApoE4 aggregation even more efficiently.^5^ In this study, we focused on studying the effect of TRN, which is a close structural homolog of TMP. TRN is an abundant amino acid produced in the liver and the central nervous system.^6^ Yet, little was known about TRN’s potential to interact with ApoE.

We first showed that TRN can inhibit ApoE4 aggregation with a similar efficiency as SPA (**Figure 4**) and that the TRN-ApoE4 interaction, similarly to that of SPA-ApoE4, specifically altered the local flexibility of ApoE4 (**Figure 5**). MD simulations revealed that TRN could destabilize dimers and inhibit aggregation, with a similar efficiency as SPA (**Figure 2A**, **Figure 3A**). The simulations suggest that these effects arise from a network of weak electrostatic interactions between the small molecules and the specific protein residues. These interactions compete with the electrostatically driven ApoE dimerization, thereby modulating its aggregation propensity (**Figure 2B-C**). Since the structure of ApoE dimers remained elusive, we employed VAMP clustering on association adaptive goal MD simulations. This analysis identified four structurally distinct dimers: (i) V-shaped, (ii) T-shaped, (iii) parallel, and (iv) anti-T-shaped. Except for the anti-T-shaped dimers, the other three were observed in the crystallographic lattice of the available ApoE N-domain crystal structures.^5^ However, the anti-T-shaped dimer offers a novel perspective on the differences between ApoE3 and ApoE4. Notably, the single mutation C112R contributes to the dimerization interface in this model, unlike the other dimer types. It is important to emphasize that the anti-T-shaped dimer model is based on MD simulations and requires experimental validation.

At the cellular level, the data collected with COs demonstrate that TRN specifically affected the biology of *ApoE4/E4* organoids, with only a minimal impact on *ApoE3/E3* organoids (**Figure 6**). Notably, TRN induced a shift in the expression of over 200 genes, bringing the gene expression profile of *ApoE4/E4* organoids closer to that of *ApoE3/E3* organoids. Notably, several of the identified differentially expressed genes were associated with neurodevelopment, autophagy pathways, or lipid metabolism, and we also confirmed this trend using MS protein analysis. Indeed, it has previously been suggested that TRN has neuroprotective effects,^46^ immunomodulatory effects,^47–49^ and the ability to activate autophagy.^50^ In the context of AD, TRN has been shown to protect neurons against excitotoxicity induced by Aβ *in vitro*,^51^ to recover spatial memory in the APP/PS1 mouse model^52^ and to improve glutamatergic activity in the brain of the 5xFAD mouse model.^53^ Our data on COs are in line with published data thus far and demonstrate that the effect of TRN is widespread. These effects appear to be linked to its direct influence on the conformation and aggregation of ApoE4, but not ApoE3 molecules. This further supports the notion that TRN, like TMP and SPA, can correct the structure, oligomerization state, aggregation propensity, and biological functions of ApoE4, shifting them closer to those of ApoE3.^5^

In summary, TRN directly interacts with ApoE4 and inhibits its aggregation, therefore promoting ApoE4’s physiological function (**Figure 7**). Moreover, it may modulate neurodegeneration through multiple mechanisms: (i) direct interaction with ApoE4 to inhibit aggregation and promote proper function, (ii) stimulation of neuron development, (iii) restoration of impaired lysosomal autophagy, and (iv) potential neuroprotective and anti-inflammatory effects.^54^ A notable similarity was observed in the effects of SPA and TRN on ApoE4 at molecular and cellular levels: these molecules induce ApoE3-like conformational changes in ApoE4 and reduce its tendency to aggregate. Dietary supplementation with TRN has already been proposed as a therapeutic strategy for addressing TRN deficiency in various pathological conditions.^55^ Our study underscores the potential of TRN as a promising therapeutic agent for treating AD and other neurodegenerative diseases associated with ApoE aggregation.

**Figure 7.**
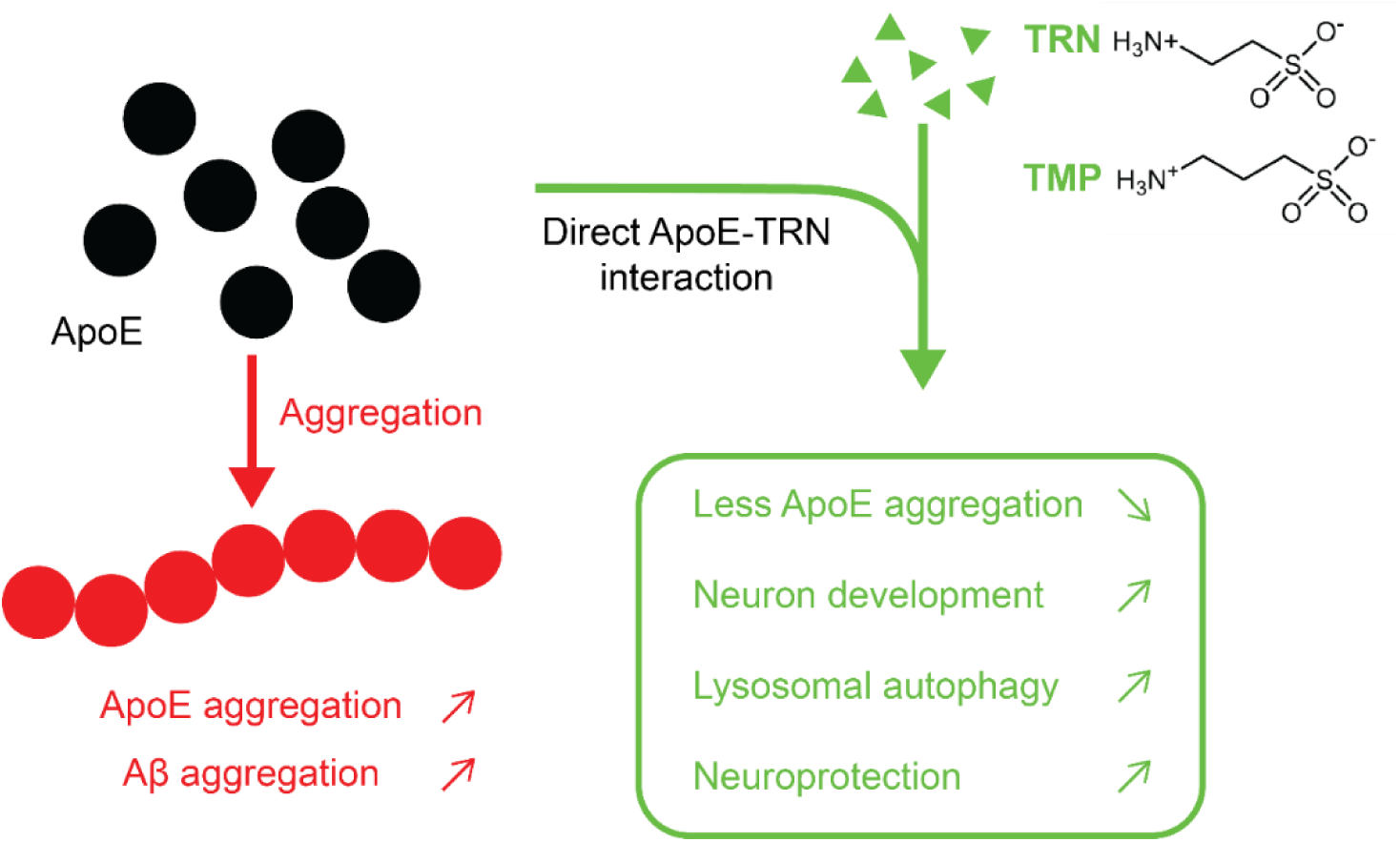
Aggregation of ApoE decreases Aβ clearance. TRN-based treatment can help in preventing ApoE aggregation through direct interaction, and in promoting neuron development, lysosomal autophagy (this study), and putative neuroprotection and anti-inflammatory properties ^54^.

## Declaration of generative AI and AI-assisted technologies in the writing process

During the preparation of this work, the authors used OpenAI’s ChatGPT to assist with language editing and stylistic improvements. After using this tool, the authors reviewed and edited the content as needed and take full responsibility for the content of the publication.

## Acknowledgements

Not applicable.

## Consent for publication

All authors have approved of the consents of this manuscript and provided consent for publication.

## Author contributions

Anthony Legrand – SLS analysis, kinetic analysis, data interpretation, manuscript writing

Katerina Amruz Cerna – cerebral organoid experiments, data interpretation

Sérgio M. Marques – computer simulations, data interpretation

Naina Verma – HDX-MS analysis, data interpretation

Jakub Kopko – VAMP analysis, data interpretation

Tereza Vanova – culturing cerebral organoids

Madhumalar Subramanian – SLS analysis

Jaroslav Bendl – statistical analysis

Tomas Henek – HDX-MS analysis

Pavel Vanacek – HDX-MS analysis, data interpretation and visualisation

Josef Kucera – HDX-MS analysis

Joan Planas-Iglesias – data interpretation

Jiri Sedmik – cultivation of cerebral organoids

Veronika Pospisilova – confocal microscopy

Petr Kouba – VAMP analysis, data interpretation

Aneta Vaskova – in vitro experiments, data interpretation

Marketa Nezvedova – targeted proteomics

Jiri Sedlar – VAMP analysis, data interpretation

Jiri Damborsky – experimental design, data interpretation, supervision

Stanislav Mazurenko – data interpretation, supervision

Martin Marek – data interpretation, supervision

Josef Sivic – VAMP analysis, data interpretation, supervision

Lenka Hernychova – HDX-MS analysis, data interpretation, supervision

David Bednar – computer simulations, data interpretation, supervision

Dasa Bohaciakova – cerebral organoids experiments, data interpretation, supervision

Zbynek Prokop – supervision, data interpretation, funding acquisition

## Data availability

The manuscript includes data available as electronic supplementary material. The mass spectrometry HDX data have been deposited to the Proteome Xchange (PX) Consortium (Deutsch et al., 2023) via Proteomics Identifications (PRIDE)^56^ partner repository with the dataset identifier PXD057427; Username; Password: 6SgqF8SmvZx5. Mass spectrometry data from targeted proteomics experiment have been deposited via Panorama Public repository, DOI: permanent link: https://panoramaweb.org/EOPVWs.url, ProteomeXchange ID: PXD059743, DOI for the data is https://doi.org/10.6069/xhy4-hj28. Sequencing data has been deposited to the GEO database at https://www.ncbi.nlm.nih.gov/geo/query/acc.cgi?acc=GSE292774, password (token): ylydkcokznqpzeh. MD simulations and VAMP analysis have been deposited to Zenodo: 10.5281/zenodo.15322679, accessible with the following link: https://zenodo.org/records/15322679?token=eyJhbGciOiJIUzUxMiJ9.eyJpZCI6IjlkNjYwNDI3LWIwNDEtNDVhNS04ZjllLTUxNjQxNjcwMjQzMCIsImRhdGEiOnt9LCJyYW5kb20iOiI4MjFhOWE2YjBkMGNmNmIxMDU2OWJmMmNlYjI4ODYzMiJ9.IEV_-vdt-F556jbNyTJU-69lfjGt9abZr2KTevPTv9ARGGRgzL5HunKdJyITHy4eTC17TTgqyVJnn9xjJwar1w. All other data supporting the findings of this study are available upon reasonable request.

