## Supporting Information for "Taurine Inhibits Apolipoprotein E4 Aggregation"

[#]Shared first authors

*Corresponding authors: Josef Sivic,; Lenka Hernychova,; David Bednar,; Dasa Bohaciakova,; Zbynek Prokop,

### Table S1. List of unique peptide transitions of analyzed proteins by LC-MS/MS

| Protein number | Gene name | Peptide sequence | ISTD? | Precursor Ion (Da) | Precursor charge | Product Ion (Da) | Product charge | Fragment ion | Retention time (min) | Collision energy (eV) |
|---|---|---|---|---|---|---|---|---|---|---|
| P10909 | *CLU* | ELDESLQVAER | FALSE | 644.8 | 2 | 802.4 | 1 | y7 | 8.85 | 21 |
| | | ELDESLQVAER | FALSE | 644.8 | 2 | 715.4 | 1 | y6 | 8.85 | 21 |
| | | ELDESLQVAER | FALSE | 644.8 | 2 | 602.3 | 1 | y5 | 8.85 | 21 |
| | | ELDESLQVAER | FALSE | 644.8 | 2 | 375.2 | 1 | y3 | 8.85 | 21 |
| | | ELDESLQVAER | TRUE | 649.8 | 2 | 812.5 | 1 | y7 | 8.85 | 21 |
| | | ELDESLQVAER | TRUE | 649.8 | 2 | 725.4 | 1 | y6 | 8.85 | 21 |
| | | ELDESLQVAER | TRUE | 649.8 | 2 | 612.3 | 1 | y5 | 8.85 | 21 |
| | | ELDESLQVAER | TRUE | 649.8 | 2 | 385.2 | 1 | y3 | 8.85 | 21 |
| P04406 | *GAPDH* | VGVNGFGR | FALSE | 403.2 | 2 | 706.4 | 1 | y7 | 4.95 | 13.5 |
| | | VGVNGFGR | FALSE | 403.2 | 2 | 649.3 | 1 | y6 | 4.95 | 13.5 |
| | | VGVNGFGR | FALSE | 403.2 | 2 | 550.3 | 1 | y5 | 4.95 | 13.5 |
| | | VGVNGFGR | TRUE | 408.2 | 2 | 716.4 | 1 | y7 | 4.95 | 13.5 |
| | | VGVNGFGR | TRUE | 408.2 | 2 | 659.3 | 1 | y6 | 4.95 | 13.5 |
| | | VGVNGFGR | TRUE | 408.2 | 2 | 560.3 | 1 | y5 | 4.95 | 13.5 |
| P11137 | *MAP2* | LINQPLPDLK | FALSE | 575.8 | 2 | 682.4 | 1 | y6 | 11.38 | 18.9 |
| | | LINQPLPDLK | FALSE | 575.8 | 2 | 472.3 | 1 | y4 | 11.38 | 18.9 |
| | | LINQPLPDLK | FALSE | 575.8 | 2 | 260.2 | 1 | y2 | 11.38 | 18.9 |
| | | LINQPLPDLK | TRUE | 579.9 | 2 | 690.4 | 1 | y6 | 11.38 | 18.9 |
| | | LINQPLPDLK | TRUE | 579.9 | 2 | 480.3 | 1 | y4 | 11.38 | 18.9 |
| | | LINQPLPDLK | TRUE | 579.9 | 2 | 268.2 | 1 | y2 | 11.38 | 18.9 |
| P48681 | *NES* | SLETEILESLK | FALSE | 631.3 | 2 | 1061.6 | 1 | y9 | 20.28 | 20.6 |
| | | SLETEILESLK | FALSE | 631.3 | 2 | 932.5 | 1 | y8 | 20.28 | 20.6 |
| | | SLETEILESLK | TRUE | 635.4 | 2 | 1069.6 | 1 | y9 | 20.28 | 20.6 |
| | | SLETEILESLK | TRUE | 635.4 | 2 | 940.5 | 1 | y8 | 20.28 | 20.6 |
| P26367 | *PAX6* | THYPDVFAR | FALSE | 369.2 | 3 | 393.2 | 1 | y3 | 7.08 | 8.5 |
| | | THYPDVFAR | FALSE | 369.2 | 3 | 246.2 | 1 | y2 | 7.08 | 8.5 |
| | | THYPDVFAR | FALSE | 369.2 | 3 | 614.3 | 1 | b5 | 7.08 | 8.5 |
| | | THYPDVFAR | TRUE | 372.5 | 3 | 403.2 | 1 | y3 | 7.08 | 8.5 |
| | | THYPDVFAR | TRUE | 372.5 | 3 | 256.2 | 1 | y2 | 7.08 | 8.5 |
| | | THYPDVFAR | TRUE | 372.5 | 3 | 614.3 | 1 | b5 | 7.08 | 8.5 |

ISTD = internal standard

**Table S2. Analytical fitting of SLS traces with an exponential burst function.** Fitting parameters, errors, and Chi-2 tests were performed with Kintek.

| Parameter | ApoE3 | ApoE4 | +SPA | +TRN |
|---|---|---|---|---|
| $A_0$ | 1936.7 | 1882.05 | 1816.92 | 1850.33 |
| Standard error ($A_0$) | 35.5942 | 152.027 | 121.892 | 198.3 |
| $A_1$ | -112910 | 5928.15 | 1575.99 | 3577.36 |
| Standard error ($A_1$) | ND | 193.436 | 149.61 | 212.087 |
| $k_{obs,1}$ | 0.016921 | 1.65588 | 1599.109 | 2.15301 |
| Standard error ($k_{obs,1}$) | ND | 0.153046 | 0.277387 | 0.314665 |
| $k_{obs,2}$ | 1912.14 | 163.609 | 398.038 | 403.909 |
| Standard error ($k_{obs,2}$) | ND | 23.6828 | 20.3908 | 19.184 |
| Δ | | | 73 ± 10 % | 40 ± 7 % |

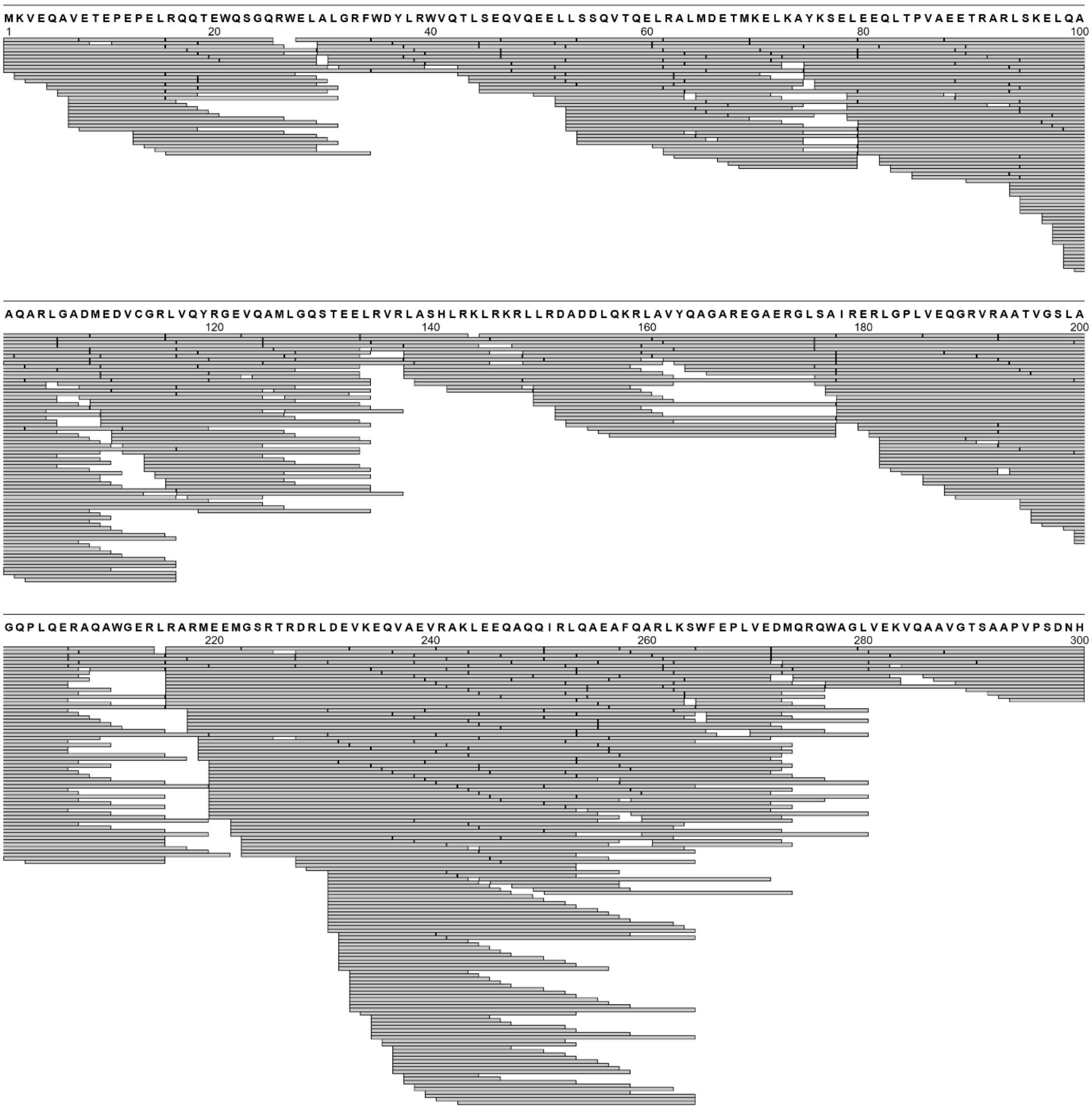

**Figure S1. Peptide coverage map of free ApoE3.** The peptides are represented by grey bars. The peptide map shows the 100% coverage of the protein sequence.

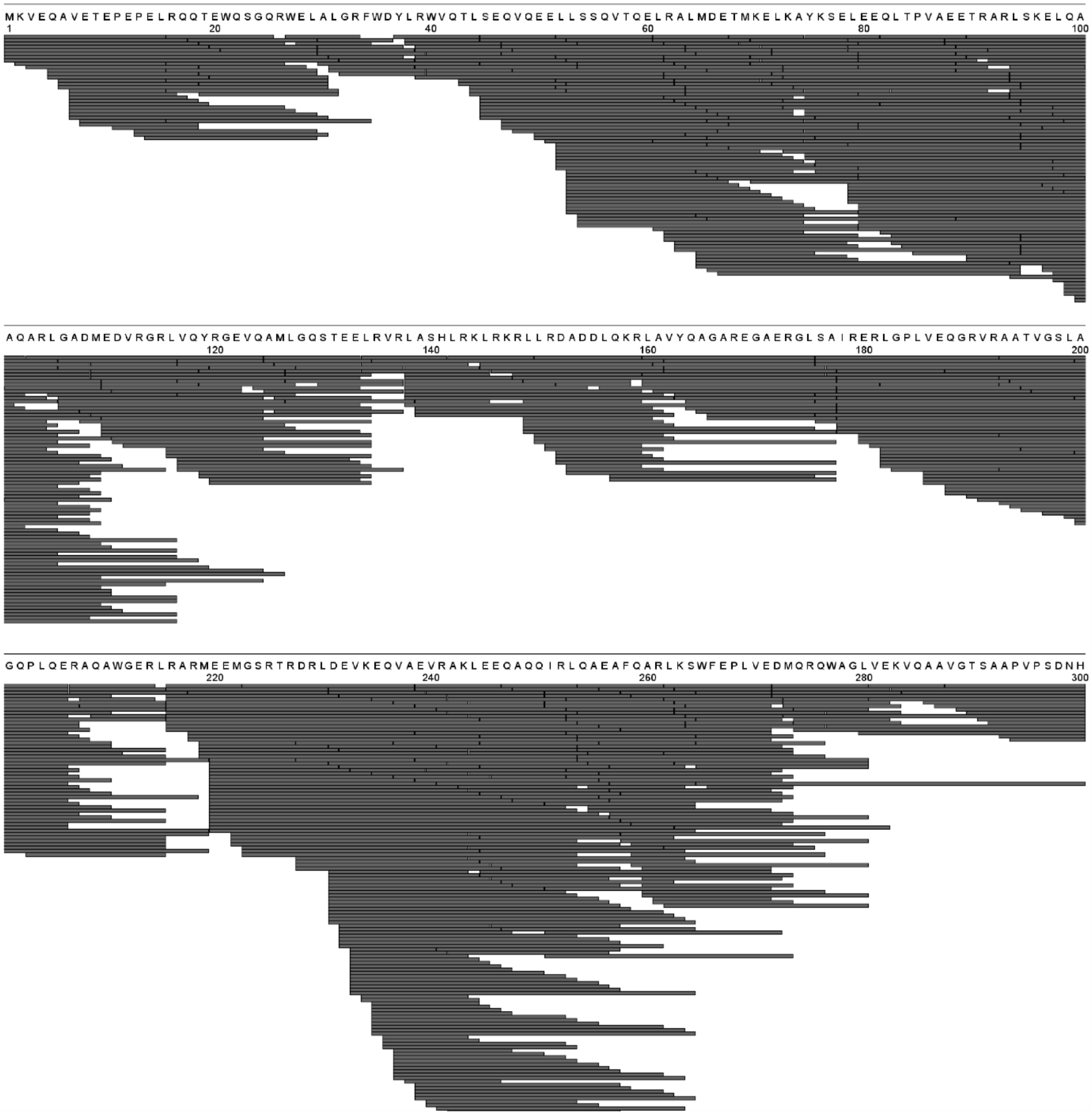

**Figure S2. Peptide coverage map of free ApoE4.** The peptides are represented by black bars. The peptide map shows the 100% coverage of the protein sequence.

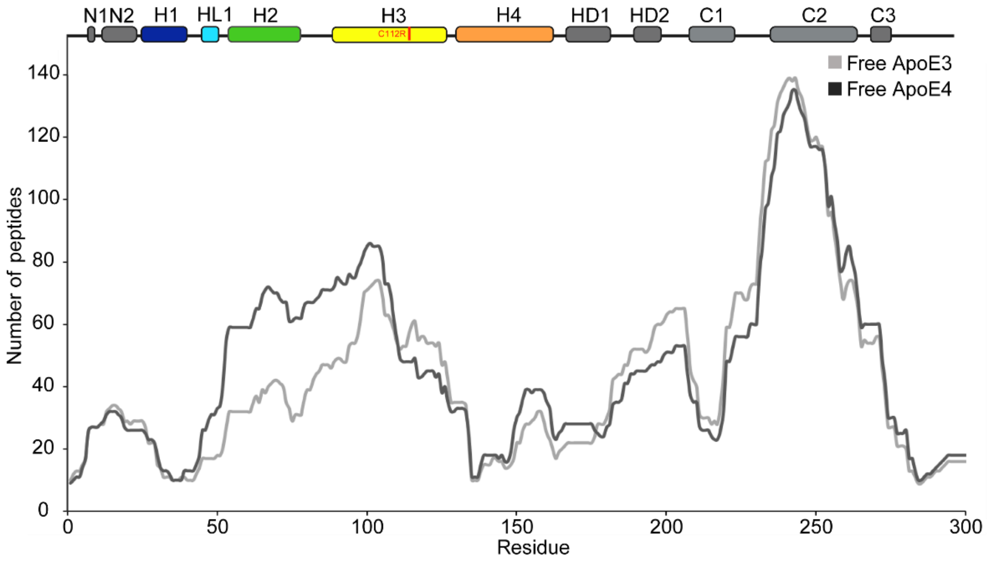

**Figure S3. Peptide redundancy plot of free ApoE3 (grey) and free ApoE4 (black).** The plot depicts the number of overlapping peptides per residue position across the protein sequence. 857 and 840 unique peptides with a redundancy score of 46.96 and 43.44 were identified for ApoE4 and ApoE3, respectively. The redundancy profile reveals regions of high peptide overlap, enabling robust HDX-MS analysis across the entire protein sequence.

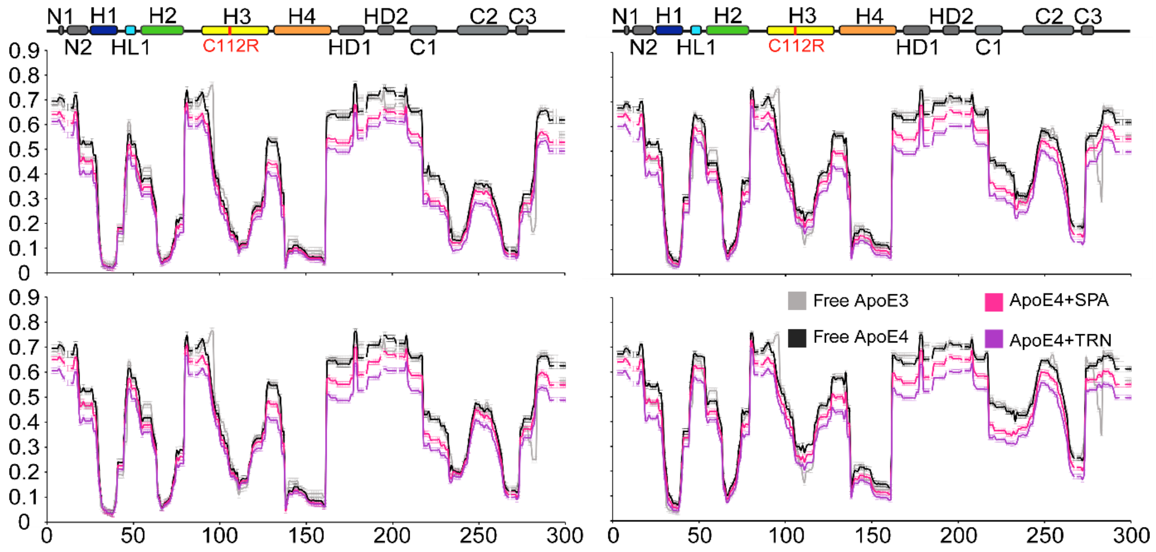

**Figure S4. Time-resolved deuterium uptake analysis of ApoE3 and ApoE4 isoforms with and without ligands.** The relative fractional uptake (RFU) is plotted as a function of residue position at deuteration time points (60, 120, 600, and 1800 sec). ApoE3 free, ApoE4 free, ApoE4 with SPA, and ApoE4 with TRN are shown in grey, black, pink, and purple, respectively. The standard error mean (SME) is depicted by the error bars.

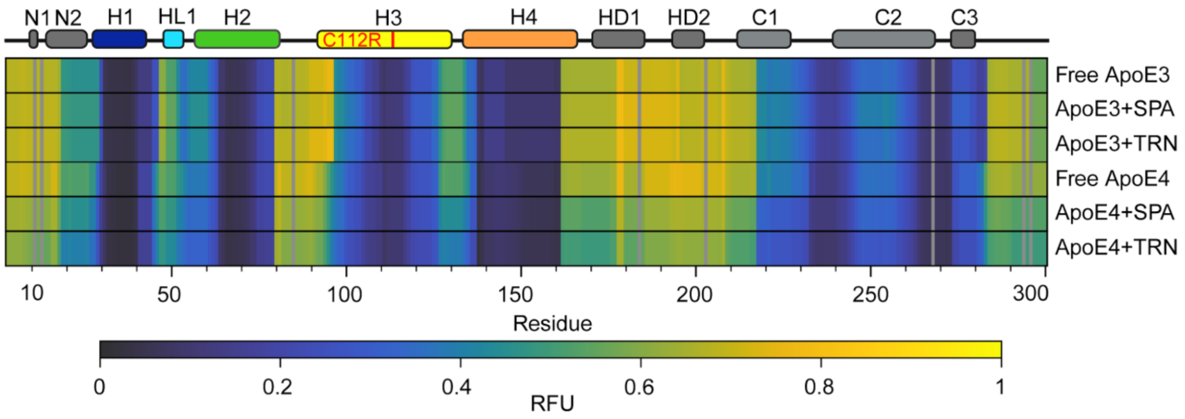

**Figure S5. Heat map showing the RFU of ApoE3/4 free and ApoE3/4 with SPA/TRN after 60 s.** The map shows that SPA and TRN have no effect on ApoE3. In contrast, with ApoE4, the RFU is significantly decreased, depicted by the green shades in the plot.

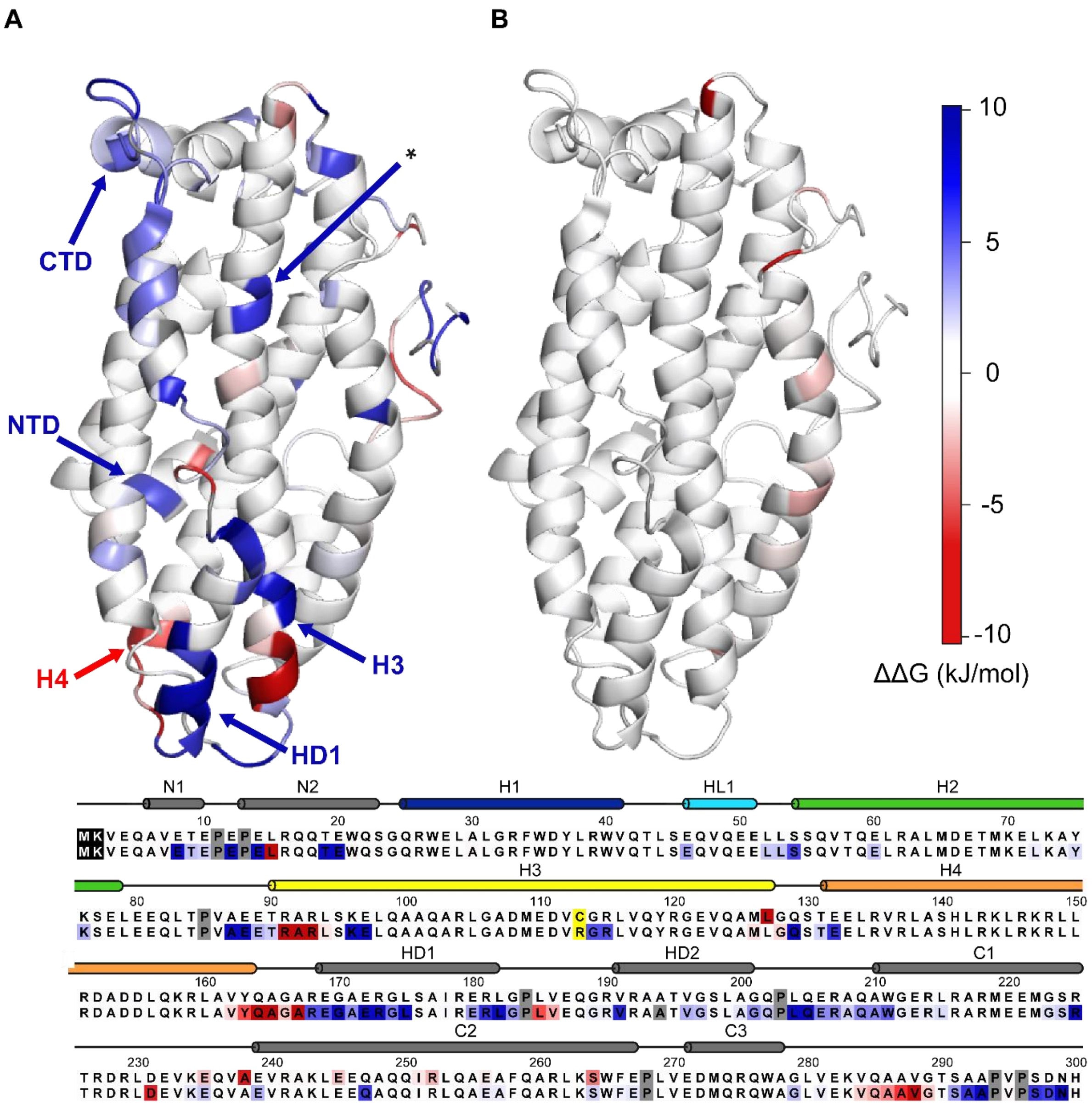

**Figure S6. Comparison of ΔΔG of ApoE3 with SPA and ApoE4 with SPA**. (A) ΔΔG values of ApoE4 with SPA, and (B) ApoE3 with SPA, mapped onto the structure of ApoE (PDB 2L7B), showing regions of altered stability. (C) ΔΔG values mapped onto the amino acid sequences of ApoE3 and ApoE4. Differences are highlighted in red and blue. Red regions indicate lower ΔΔG values, suggesting increased amide hydrogen (H) to deuterium (D) exchange compared to the free protein, while blue regions indicate reduced exchange. The mutation site (C112R) is highlighted in yellow with an asterisk, and the first two residues, excluded from pyHDX analysis, are shown in black. Proline residues, non-exchangeable in HDX-MS, are indicated in grey.

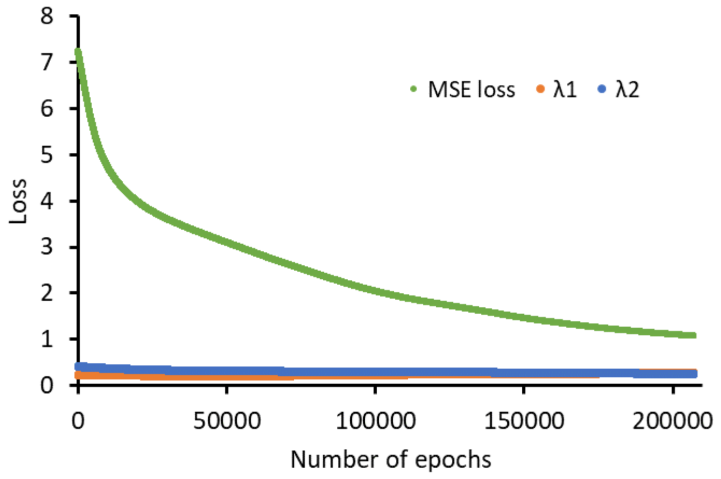

**Figure S7. Fit losses (Lagrangian value) per epoch.** Mean square error (MSE) loss is depicted by green colour with total loss of 1.59. The regularizer 1 (λ1) and regularizer 2 (λ2) are depicted by orange and blue colours, respectively. The regularization loss is 0.52 (32.6%).

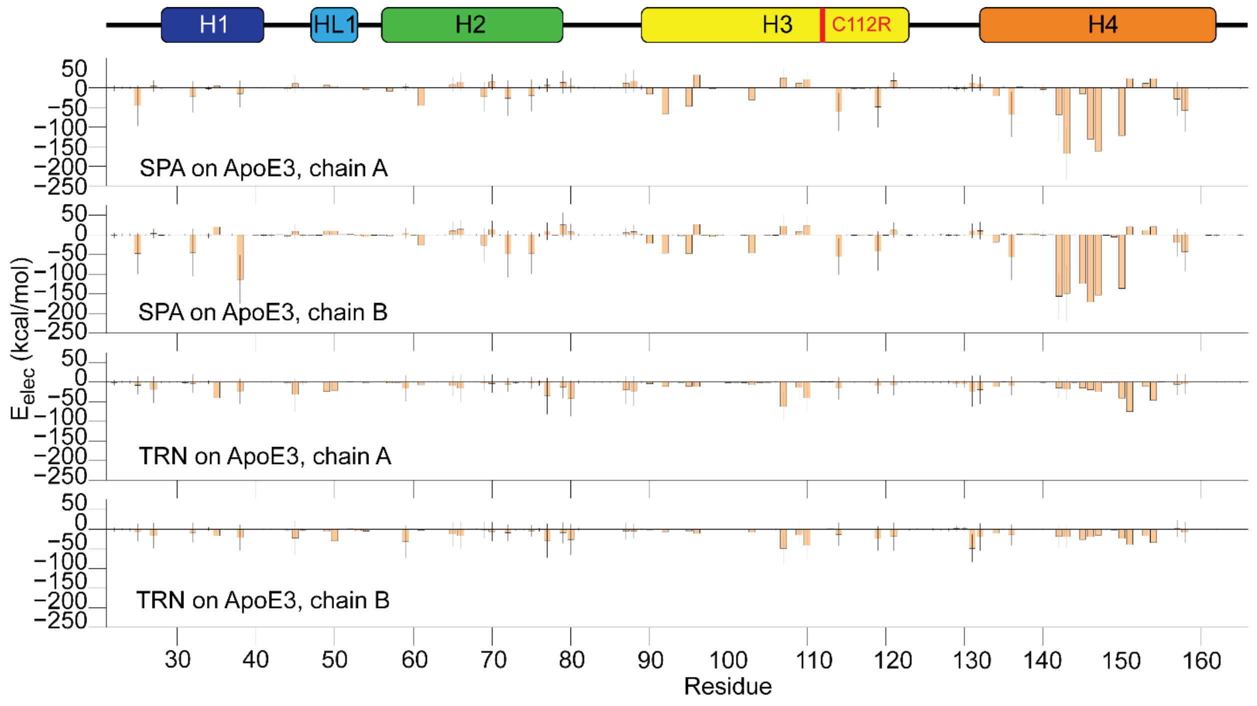

**Figure S8. Electrostatic component of LIE of ApoE3 T-shaped dimers in the presence of SPA or TRN.** Stick figure of ApoE3 showing the secondary elements and the position of C112R mutation.

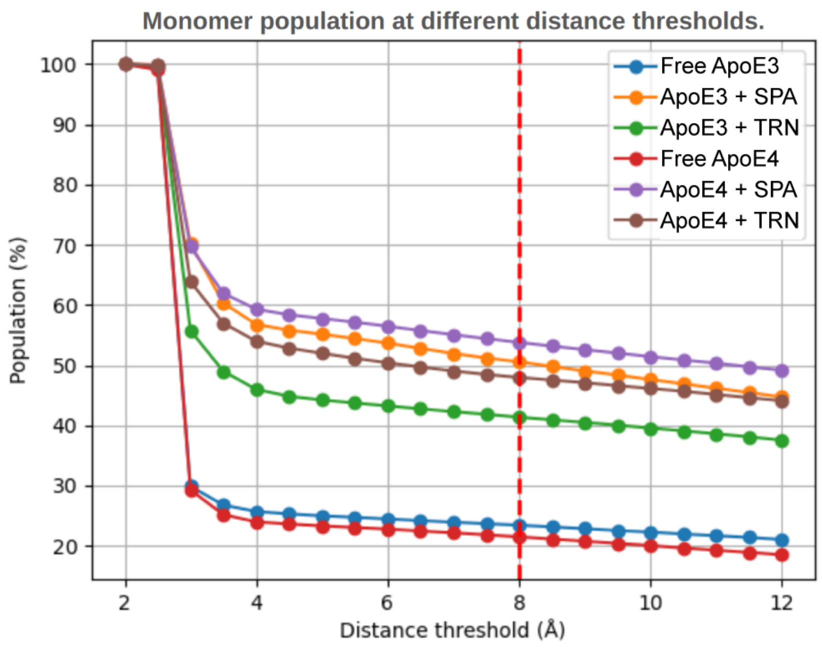

**Figure S9. Computed number of frames representing monomers.** The threshold is robust across different distance thresholds. Ultimately, a threshold of 8 Å was chosen (marked with the vertical dashed red line). The X-axis represents the distance threshold used for counting monomeric frames. The Y-axis represents the population of monomeric frames expressed as a fraction of the total number of frames in a given simulation. The total number of monomeric frames remains relatively stable across varying thresholds, and the relative ranking of systems by their monomeric frame count stays consistent. This consistency demonstrates that our observations are independent of the chosen threshold.

**Table S3. Frame classification of the association goal simulations.** The assignment rules are as follows: when the distance between chains exceeds 8 Å, the structure is classified as a monomer. If the symmetrized RMSD to all dimer forms (T, V, anti-T, parallel) exceeds the "quality threshold" of 6 Å, the structure is classified as "Other." Otherwise, the structure is assigned to the type of dimer with the lowest RMSD. Notable observations (in bold) include the prevalence of the parallel dimer in the free ApoE4 system (7.6%), which is significantly reduced in the presence of TRN (1.7%) and SPA (0.0%) and is absent in ApoE3 systems. Additionally, the existence of the anti-T-shape dimer is observed in the free ApoE4 system (4.8%). We can observe the highest fraction of dissociated frames in the ApoE3+SPA (50.5%) and ApoE4+SPA (53.8%) systems. The impact of the RMSD threshold value is assessed in Figure S8.

| System | Dimer Parallel | Dimer V-shape | Dimer Anti-T-shape | Dimer T-shape | Other | Monomer |
|---|---|---|---|---|---|---|
| ApoE3 | 0.5% | 0.3% | 0.0% | 3.5% | 72.3% | 23.4% |
| ApoE3 + TRN | 0.0% | 0.3% | 0.0% | 0.5% | 57.7% | 41.4% |
| ApoE3 + SPA | 0.1% | 0.0% | 0.0% | 0.0% | 49.4% | **50.5%** |
| ApoE4 | **7.6%** | 0.9% | **4.8%** | 1.1% | 64.2% | 21.5% |
| ApoE4 + TRN | **1.7%** | 0.0% | 0.0% | 0.0% | 50.3% | 48.0% |
| ApoE4 + SPA | **0.0%** | 0.0% | 0.0% | 0.1% | 46.1% | **53.8%** |

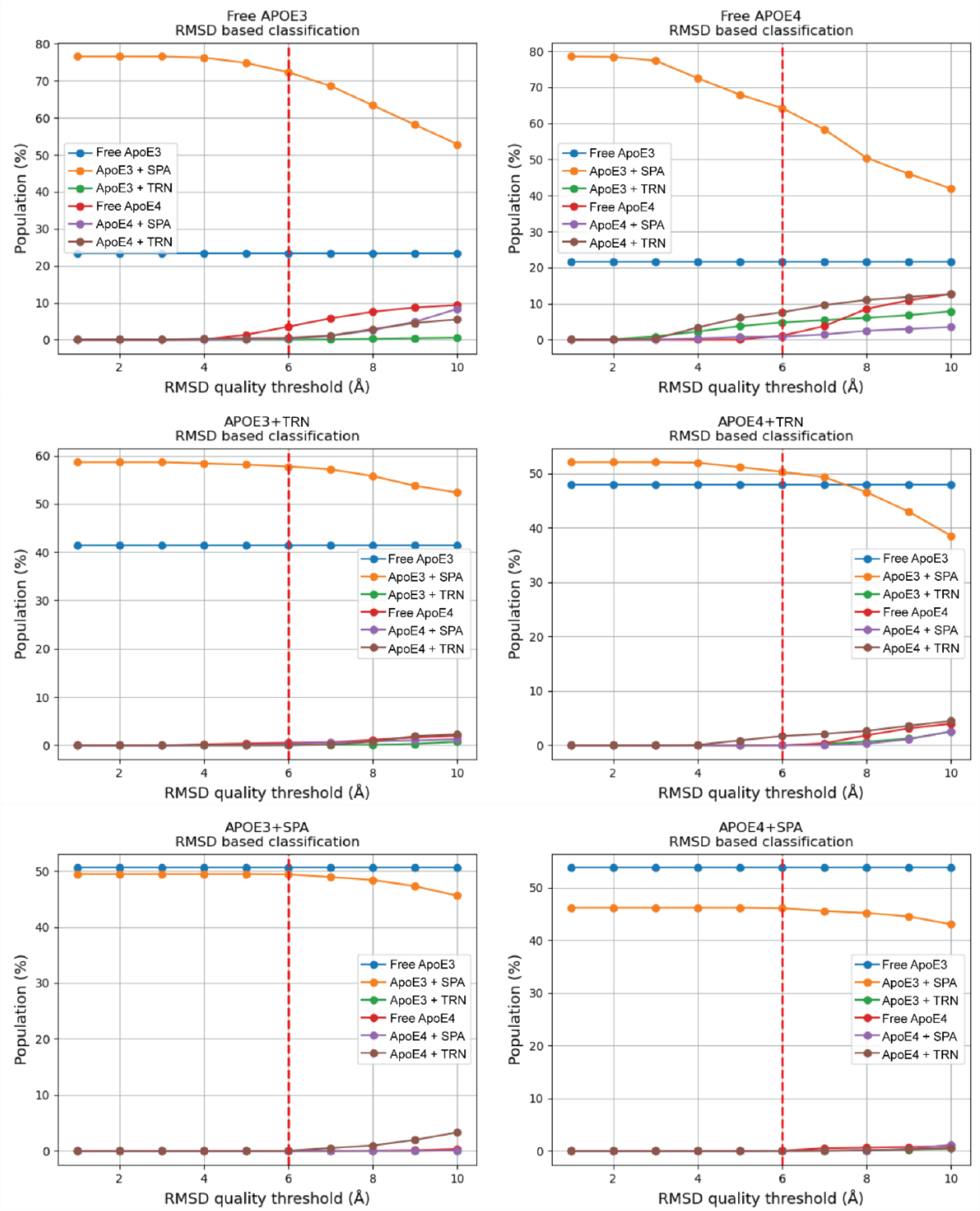

**Figure S10. Impact of RMSD quality threshold on frame classification in Table S3.** Populations of each frame class as a function of the RMSD quality threshold above which a non-“monomer“ frames is classified as “Other“. Naturally, increasing the RMSD quality threshold increases the populations of classes corresponding to reference structures, but the observations we focused on are robust to the choice of “quality threshold”, namely parallel dimer being the dominant dimer of free ApoE4 and anti-T-shape dimer existing only in ApoE4. Blue: monomers. Orange: others. Green: anti-T-shape dimer. Red: T-shaped dimer. Purple: V-shape. Brown: parallel dimer.

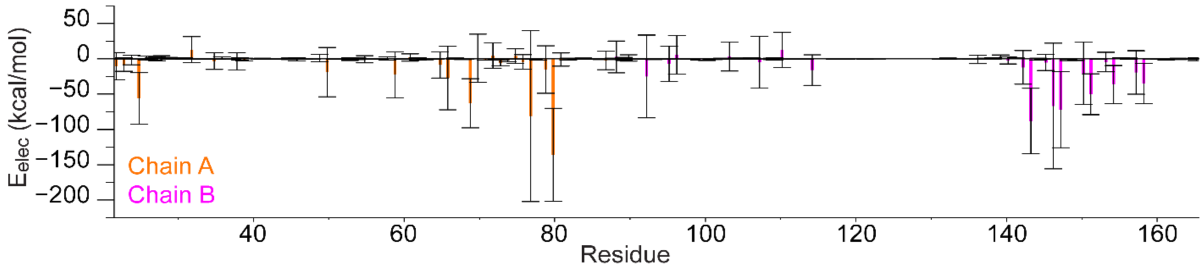

**Figure S11. Inter-chain electrostatic component of LIE of ApoE4 parallel dimers.** Evaluated on 1000 random frames from the VAMP cluster representing free ApoE4 in parallel dimer conformation (yellow cluster in Figure 5B), and plotted for each residue and each chain. Orange: chain A. Purple: chain B.

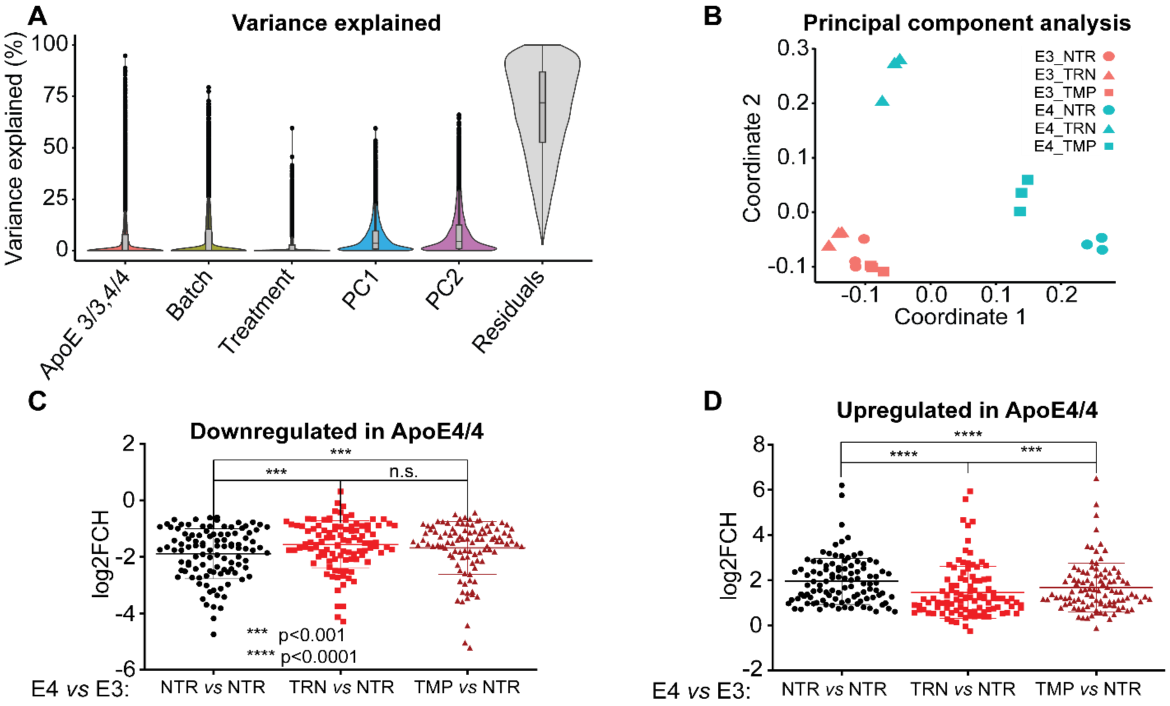

**Figure S12. Effects of TMP and TRN treatment on cerebral organoids on the transcriptome.** (A) Variance explained by biological and technical covariates calculated on normalised read counts. (B) Multidimensional scaling including samples treated with TRN or TMP. (C,D) LogFoldChange expression of genes differentially expressed in between ApoE4/ApoE3 genotypes (p-adj<0.1, abs(log2FCH)>0.6; n=206) and their shift in expression after TRN or TMP treatment (C) Genes down-regulated in ApoE4 compared to ApoE3 and shift of their expression toward the ApoE3 upon the TRN/TMP treatment (p<0.001; n=104). (D) Genes up-regulated in ApoE4 compared to ApoE3 and shift of their expression toward the ApoE3 upon the TRN treatment (p<0.001; n=102).

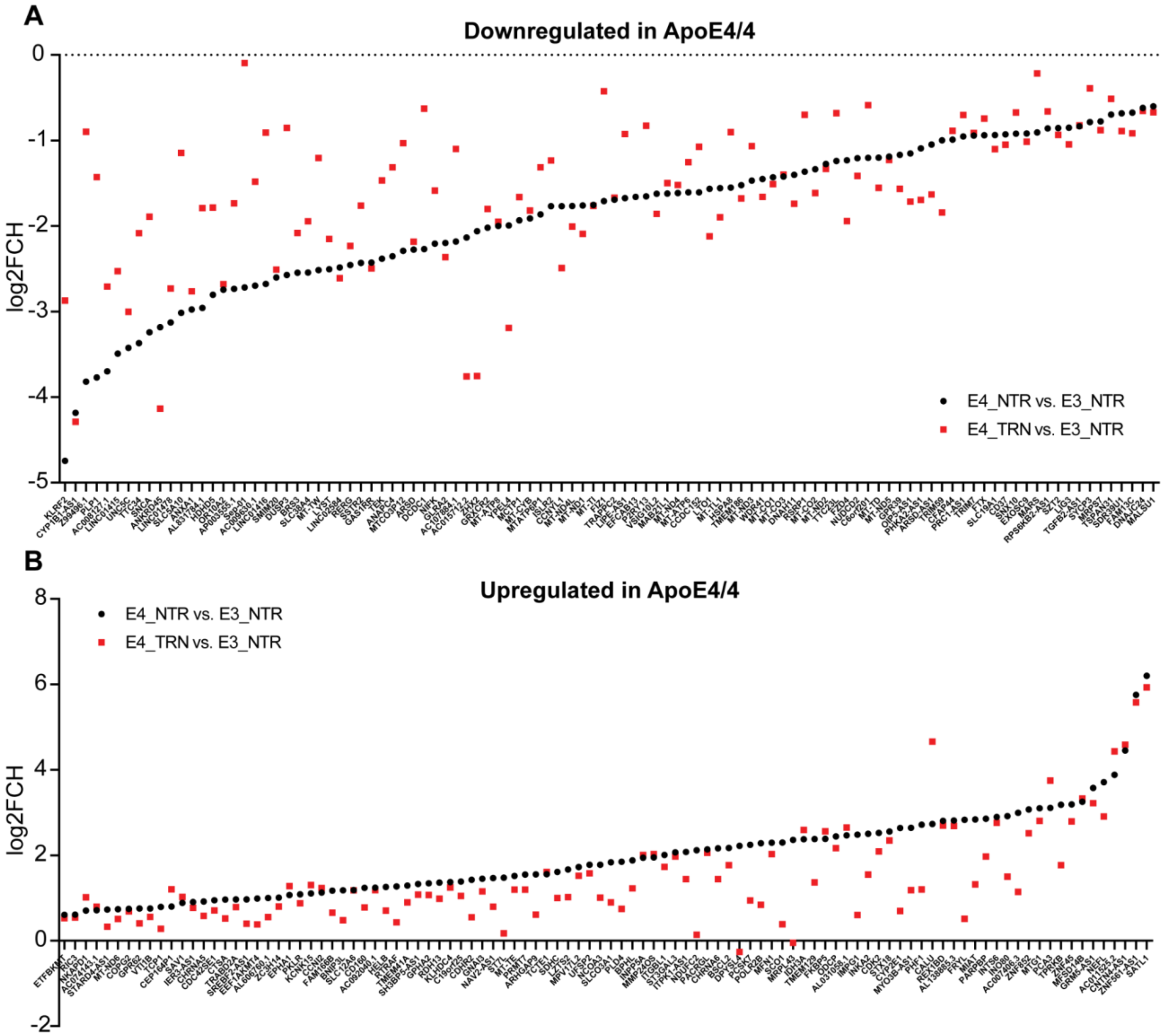

**Figure S13. Detailed visualisation of log(2FCH) expression of differentially expressed genes.** Comparison between ApoE4 and ApoE3 genotypes. (p-adj<0.1, abs(log2FCH)>0.6; n=206) and their shift in expression after TRN or TMP treatment. (A) Down- and (B) up-regulated genes in ApoE4/4 COs as a function of TRN-treatment.

**Table S4. List of log(2FCH) expression of differentially expressed genes between ApoE4 and ApoE3 genotypes**

See Additional file 1

**Table S5. Analysis of differential expression.** As described in the material and methods section for comparisons of E4_NTR vs. E3_NTR (list 1), E4_TRN vs. E3_NTR (list 2) COs.

See Additional file 2

**Table S6. Differentially expressed genes in ApoE4 COs upon TRN treatment.**

| Biological process | ENSG | Gene name | logFC_E4.NTR_E3.NTR | logFC_E4.TAU_E3.NTR |
|---|---|---|---|---|
| Lipid metabolism | ENSG00000130649 | CYP2E1 | 2.636183352 | 0.69619 |
| Lipid metabolism | ENSG00000168000 | BSCL2 | 2.169327024 | 1.77141 |
| Lipid metabolism | ENSG00000126091 | ST3GAL3 | 2.068173235 | 1.97091 |
| Lipid metabolism | ENSG00000166428 | PLD4 | 1.846800792 | 0.74373 |
| Lipid metabolism | ENSG00000176463 | SLCO3A1 | 1.833952275 | 0.89847 |
| Lipid metabolism | ENSG00000160439 | RDH13 | 1.358262423 | 0.98424 |
| Lipid metabolism | ENSG00000139160 | ETFBKMT | 0.605337917 | 0.52994 |
| Lipid metabolism | ENSG00000139160 | ETFBKMT | 0.605337917 | 0.52994 |
| Lipid metabolism | ENSG00000120156 | TEK | -2.381856524 | -1.46684 |
| Lipid metabolism | ENSG00000069998 | HDHD5 | -2.804115384 | -1.78389 |

| | | | | |
|---|---|---|---|---|
| Lipid metabolism | ENSG00000135046 | ANXA1 | -2.976416751 | -2.76269 |
| Lipid metabolism | ENSG00000145335 | SNCA | -3.241299542 | -1.89066 |
| Lipid metabolism | ENSG00000123560 | PLP1 | -3.769617504 | -1.42849 |
| Neurodevelopment | ENSG00000277586 | NEFL | 3.70865695 | 2.90701 |
| Neurodevelopment | ENSG00000075539 | FRYL | 2.828738767 | 0.51367 |
| Neurodevelopment | ENSG00000160439 | RDH13 | 1.358262423 | 0.98424 |
| Neurodevelopment | ENSG00000167178 | ISLR2 | -1.768403912 | -1.23310 |
| Neurodevelopment | ENSG00000204928 | GRXCR2 | -2.020544831 | -1.80164 |
| Neurodevelopment | ENSG00000135046 | ANXA1 | -2.976416751 | -2.76269 |
| Neurodevelopment | ENSG00000182168 | UNC5C | -3.423849378 | -3.00236 |
| Neurodevelopment | ENSG00000123560 | PLP1 | -3.769617504 | -1.42849 |
| Autophagy | ENSG00000171109 | MFN1 | 2.295061221 | 2.02837 |
| Autophagy | ENSG00000065135 | GNAI3 | 1.448630028 | 1.15606 |
| Autophagy | ENSG00000104765 | BNIP3L | 1.179430288 | 0.47754 |
| Autophagy | ENSG00000064601 | CTSA | 0.963330891 | 0.51948 |
| Autophagy | ENSG00000109971 | HSPA8 | -1.55198484 | -0.90219 |
| Autophagy | ENSG00000145335 | SNCA | -3.241299542 | -1.89066 |
